# Reprogramming CAR-T cell function through synthetic cytokine receptors suppresses metastatic progression of TNBC

**DOI:** 10.64898/2026.10.22.760671

**Authors:** Eban A. Hanna, Alice Amitrano, Andre Forjaz, Zhiliang Bai, Chanhong Min, Yoseph Dance, Immanuel Etoh, Bryce Lloyd, Xinyun Jiang, Vasco Queiroga, Praful Nair, Victoria Duarte, Saurabh Joshi, Konstantinos Konstantopoulos, Jude Phillip, Rong Fan, Pei-Hsun Wu, Denis Wirtz

## Abstract

Patient T cells modified with chimeric antigen receptors (CARs) have shown remarkable therapeutic success in previously incurable cancers. However, despite its success in hematological malignancies, CAR-T therapy has shown only modest efficacy in solid tumors. The medical field has focused on developing new molecular designs that enhance CAR-T persistence, prevent exhaustion in the tumor microenvironment (TME), and increase the specificity of the CAR-T cell to various tumor antigens. However, much less effort has been placed on enhancing CAR-T cells’ invasion into and migration through the stromal matrix of the TME, where they can elicit tumor-directed cytotoxicity. Here, we engineered CAR-T cells with synthetic cytokine receptors targeting inflammatory cytokines produced by the cells themselves, which increase their speed through collagen I matrices. We show that this enhanced motility is driven by increased CAR-T cell translocation through confined microchannels via cellular deformation, together with elevated migratory speeds in proximity to neighboring cells, suggesting improved responsiveness to local cues. These migration-enhanced CAR-T cells significantly reduce both primary tumor size and distant organ metastasis in triple-negative breast cancer (TNBC) mouse models compared to standard CAR-T cells. These results support molecular design strategies that exploit mechanisms to improve cellular therapies for metastatic solid tumors.

## Introduction

CAR-T cell therapy has resulted in remarkable response rates and remission in patients with hematological malignancies^1,2^. However, their efficacy in solid tumors remains disappointing^3^, limited by several microenvironmental factors. Cancer cells secrete anti-inflammatory cytokines such as TGF-β and IL-10 into the tumor microenvironment (TME), downregulating active immune responses^4,5^, while differentiating anti-tumor effector cells into more immunosuppressive and pro-tumoral phenotypes^6^. Malignant cells also express immune checkpoints, such as PD-L1, promoting exhaustion and cellular dysfunction of activated cytotoxic lymphocytes during cell-to-cell contact with their cognate ligands^7^. However, prior to facing these molecular barriers, CAR-T cells must first effectively traffic to the tumor cells. Improvements in CAR constructs are futile if the resulting CAR-T cells cannot invade and infiltrate the tumor. Due to the presence of thick stromal extracellular matrices (ECM) in the TME, post-IV injection, a large fraction of these CAR-T cells do not penetrate and migrate effectively through the stromal boundary of the tumor mass^8–10^, contributing to the poor efficacy of current cellular therapies in solid tumors. Increasing the therapeutic dose to “force” infiltration into the tumor is often not an appropriate option due to increased risk of off-target effects.

Breast cancer is an aggressive form of epithelial adenocarcinoma, accounting for 32% of new cancer cases in women^11^. Cancer-associated fibroblasts (CAFs) make up a significant portion of the breast cancer TME, impeding T cell infiltration through collagen deposition^12^. Triple-negative breast cancer (TNBC), a more aggressive subtype of breast cancer which is characterized by the absence of estrogen, progesterone, and human epidermal growth factor receptors, features an even thicker, denser, and more fibrotic stromal matrix compared to other breast cancer subtypes^13^. TNBC tumors maintain lower CD8+ T cell infiltration compared to non-TNBC breast cancer, due in part to the increased fibrotic nature of the surrounding TME^14^. Patients with TNBC tumors with high infiltration of CD8+T cells exhibit a better prognosis compared to patients with poor T cell infiltration ^15^.

Although collagen deposition contributes to the overall inefficacy of CAR-T cell therapy in treating solid tumors, limited efforts have focused on understanding the biophysical mechanisms underlying T cell migration and how these mechanistic insights could be used to design new CAR-T therapies that infiltrate the tumor microenvironment (TME) more effectively. While immunological mechanisms are essential for evaluating therapeutic efficacy, they provide an incomplete view of CAR-T cell function. Critically, they overlook the physical constraints and behaviors that ultimately govern whether CAR-T cells can reach, infiltrate, and engage tumor cells. Among these, cell migration is not merely a supporting feature but a fundamental determinant of therapeutic success. In the context of solid tumors, in particular, the ability of CAR-T cells to navigate complex, confined microenvironments is a key physical process that remains underexplored, despite its direct impact on treatment outcomes^16,17^.

To combat the molecular elements in the TNBC microenvironment that result in poor infiltration and efficacy of cytotoxic T cells^11,20,22,31^ we studied a phenomenon in which T cells move faster when seeded at higher cell densities due to the secretion of IL-5, IL-8, and TNF-α. When added to T cells, these cytokines, which are expressed at low levels by T cells themselves, push them into a high-migratory phenotype. Taking advantage of this concept, we engineered synthetic cytokine receptors that lock standard mesothelin-targeting M5 CAR-T cells into this high-migratory phenotype^18^. CAR-T cell migration was evaluated using 3D collagen gel assays and microfluidic devices that model confined microenvironments, enabling identification of the physical and molecular mechanisms underlying enhanced motility. Lastly, these modified CAR-T cells were compared to standard CAR-T cells across multiple in vitro immunological assays and in vivo models of metastasis to quantify tumor burden at the primary site and in the lungs.

## Results

### Harnessing cell density-based cytokine signaling to engineer synthetic cytokine receptors that enhance CAR-T cell migration

Following our observation that cancer cells migrate more rapidly at higher seeding densities compared to lower densities^19^we investigated whether this phenomenon extends to T cells. Non-engineered T cells were seeded at low, medium, and high densities (10K, 50K, and 100K cells/mL, respectively). Similar to cancer cells, T cells exhibited a density-dependent increase in motility, with higher cellular densities corresponding to increased migration (Fig. 1a).

**Figure 1.**
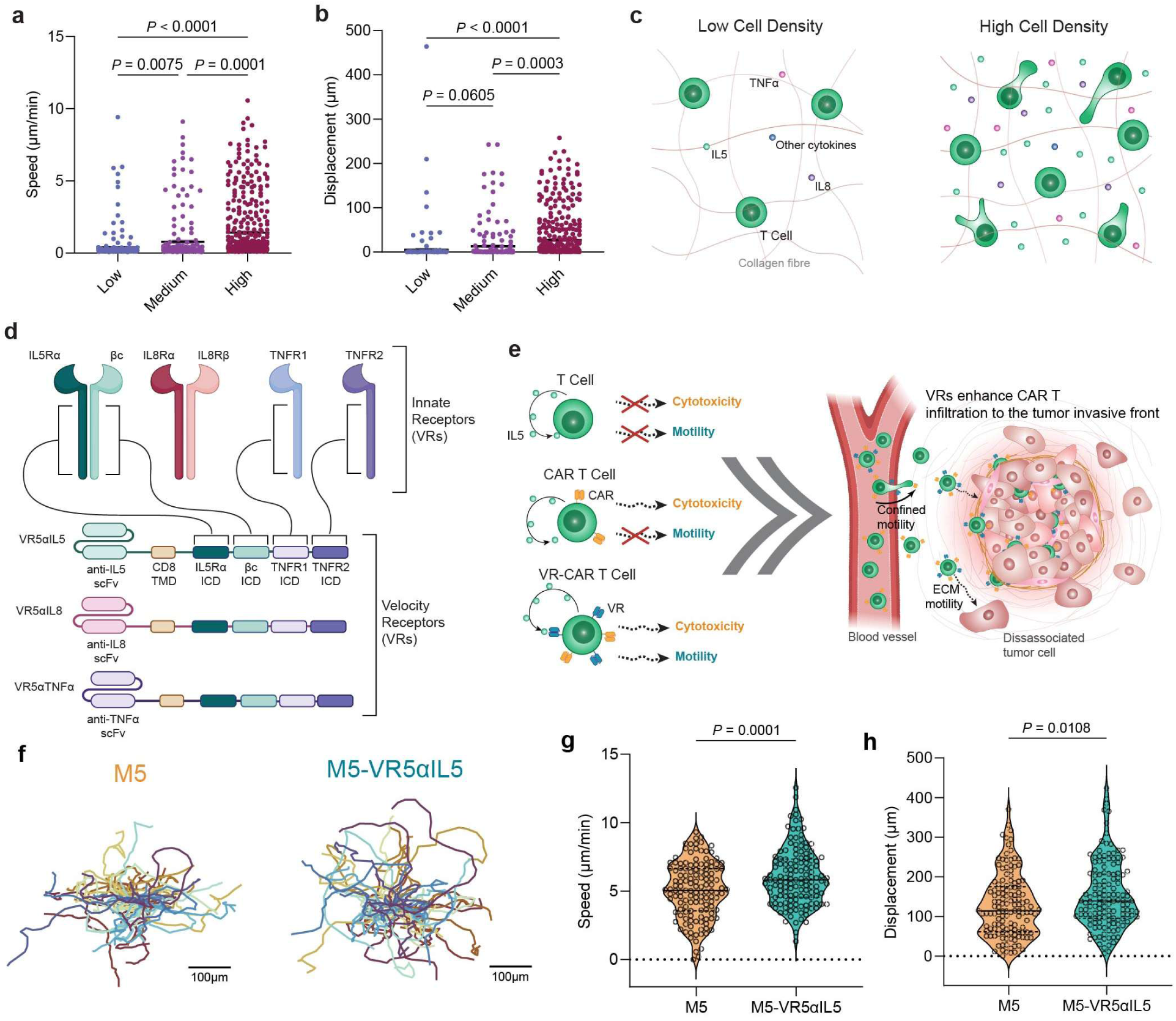
Synthetic cytokine receptors designed to signal through native IL-5 and TNFα intracellular domains promote enhanced migration in CAR-T cells. (a,b) Non-transduced T cells seeded in collagen I gels show increased speed and displacement at higher cell densities. Data plotted as mean ± SEM, N=2 biological replicates, n=3 technical replicates (>223 cells per condition). One-way ANOVA with Games-Howell’s multiple comparisons test was used to assess statistical significance. (c) Schematic illustration of non-transduced T cells seeded at low (10,000 cells/mL), medium (50,000 cells/mL), and high (100,000 cells/mL). T cells seeded at high density secrete elevated amounts of IL-5, IL-8, and TNFα compared to lower densities. (d) Diagram of mesothelin-targeting M5CAR and synthetic velocity receptor (VR) constructs. Synthetic receptors are derived from the signaling components of the native IL-5 and TNFα receptors, fused together with an scFv targeting either IL-5, IL-8, or TNFα. (e) VR augments CAR-T cells with improved motility features, allowing them to better infiltrate solid tumors to stop cancer growth and dissemination. (f) Twenty randomly selected trajectories of M5 and M5-VR5αIL5 CAR-T cells in the collagen matrices. Scale bar = 100 µm. (g, h) Speeds and displacements of M5 and M5-VR5αIL5 CAR-T cells during 1-h time-lapse videos collected at 2-min intervals. Violin plots display the median with interquartile range (IQR), N=3 biological replicates, n=3 technical replicates (>132 cells per condition). Welch’s t-test was used to assess statistical significance.

Previous studies have shown that elevated T cell density is associated with increased secretion of inflammatory cytokines, including IL-5, IL-8, and TNFα, which correlate with enhanced migratory behavior under these conditions^18^ (Fig. 1b). Motivated by this relationship between cytokine-rich microenvironments and increased motility, we engineered CAR-T cells to express a synthetic cytokine receptor, termed “velocity receptor (VR)”, designed to sense and respond to these autocrine and paracrine signals. This receptor incorporates intracellular signaling domains derived from the IL-5 receptor complex (IL5Rα and βc) as well as TNFR1 and TNFR2, and is equipped with scFvs targeting IL-5, IL-8, or TNFα, enabling CAR-T cells to sense and engage cytokine signals enriched in high-density environments (Fig. 1d and Supplementary Fig. 1a).

We hypothesized that enhancing CAR-T cell sensitivity to density-associated cytokine cues would improve their ability to navigate confined tumor microenvironments, thereby increasing tumor infiltration and enabling more effective clearance of both static and disseminating tumor cells (Fig. 1e). To test this hypothesis, T cells were first transduced with the velocity receptor and subsequently transduced with a standard mesothelin-targeted CAR previously evaluated in clinical trials for mesothelin-expressing solid tumors^20^ (Supplementary Fig. 1b-c). Consistent with our hypothesis, CAR-T cells expressing the VR exhibited increased track speed and displacement within collagen I matrices (Fig. 1f–h), indicating improved migratory performance in physiologically relevant confined environments. These results support the rationale for engineering synthetic cytokine receptors to enhance CAR-T cell motility and infiltration, thereby improving efficacy against diseased cells protected by dense extracellular matrices.

### Synthetic cytokine receptor signaling induces a distinct motility phenotype in CAR-T cells migrating within 3D collagen gels

To fully characterize the motility of CAR-T cells possessing the VR, we seeded modified CAR-T cells in collagen I gels and manually tracked the trajectory of single cells over an hour using brightfield microscopy. Given that IL-5 was found to be secreted in greater quantities compared to the other inflammatory cytokine observed^18^, we chose to use the M5-VR5αIL5 as a proof of concept for the advanced motility analyses. After verifying our hypothesis that speed would be improved (Fig. 1g-h), we sought to determine other motility-associated parameters that would explain the results observed. It has been shown that single-cell migration is a result of more than speed or displacement alone^21^. For example, metrics such as tortuosity quantify the efficiency with which a cell moves from point A to point B, distinguishing between directed, near-linear trajectories and more exploratory Lévy-flight-like movement patterns. We first took a time-based approach to determine how much time these cells spend in a “motile state”, defined as a period of time between frames in which a cell moves a distance greater than its radius (assumed to be 5 microns for T cells) and the speed of their state once within it (Fig. 2a). M5-VR5αIL5 CAR-T cells were found to spend more time in motile states, while maintaining higher speeds within them (Fig. 2, b-d).

**Figure 2.**
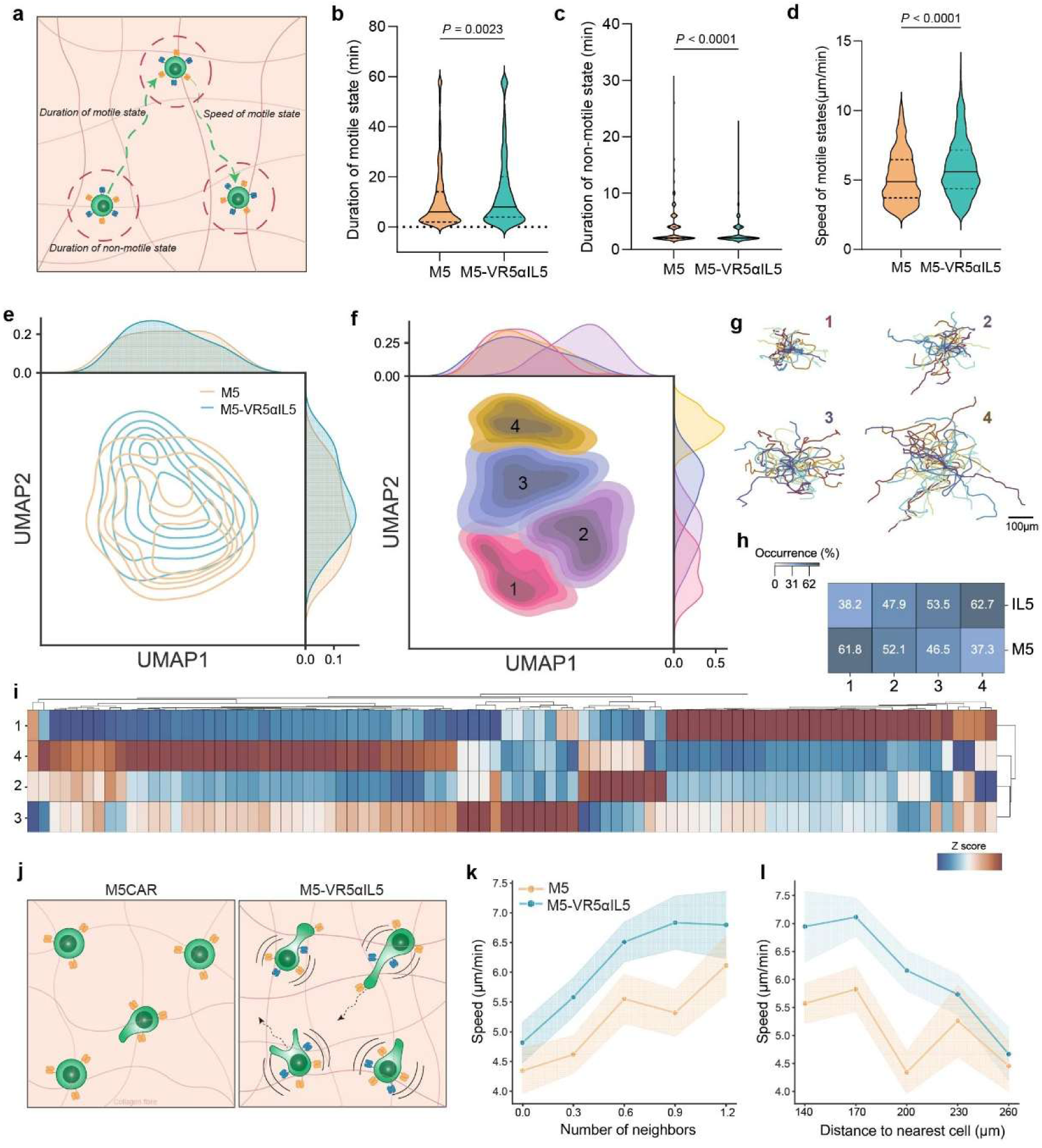
Synthetic cytokine receptors enhance multiple features of CAR-T cell motility through proximity-dependent paracrine signaling. (a) A stop-and-go motion analysis of CAR-T cells was performed to compare specific characteristics of motile states. (b-d) VR5αIL5 spend more time in a motile state and have higher speeds during motile states. Violin plots display the median with interquartile range (IQR). Welch’s t-test was used to assess statistical significance. (e,f) Contour overlay of UMAP spaces based on 88 motility features across both conditions. Clustering was determined using k-means. (g) Twenty randomly selected trajectories of each UMAP cluster. Scale bar = 100 µm. (h) Proportion of each condition in every cluster. (i) Heat map showing Z-score expression of different motility features in each cluster of the UMAP space. Positive Z-scores indicate upregulation in that specific motility feature compared to the other clusters. (j-l) Nearest-neighbor analysis to determine effects of paracrine signaling and cell-to-cell distance on motility. VR5αIL5 shows increased speeds when closer to other cells and converges with M5 speeds at larger distances from each other. (a-m) N = 3 biological replicates, M5: n = 127, M5-VR5αIL5: n = 135 cells.

Using a multidimensional motility analysis, more than 80 motility features were analyzed to extract insights into the VR-CAR-T’s motility pattern. Most features were strongly co-expressed in M5-VR5αIL5 except for display autocorr_y_1, net distance_y, angle_gini, angle_cov, and inst_angle_form factor (Supplementary Fig. 2a). Through differential analysis, we show that most features that were upregulated in the IL-5-targeting condition correspond to faster cell movement, as shown through features such as MSD, speed, and distance traveled. Features such as displacement_gini and displacement covariance, however, were found to be downregulated in M5-VR5αIL5 compared to M5 CAR alone (Supplementary Fig. 2b). We then constructed a UMAP space based on all motility features to identify clusters specific to M5-VR5αIL5 that were upregulated in those corresponding to increased motility. Clusters 3 and 4 possessed cells that moved long distances, as shown through their related trajectories, as well as the highest occurrence of the M5-VR5αIL5 condition, while clusters 1 and 2 were mostly composed of the M5 condition (Fig. 2e-h and Supplementary Fig. 2c). These same clusters, possessing a higher occurrence of M5-VR5αIL5 cells, had the highest Z-score for features mostly associated with raw speed, such as MSD, speed, and displacement, proving that the M5-VR5αIL5 condition did have a higher percentage of high motility cells compared to the M5 CAR condition alone (Fig. 2i and Supplementary Fig. 2d). Lastly, a nearest neighbor’s analysis reveals that the VR-CAR-T cell’s enhanced migration was dependent on their proximity to nearby cells (Fig. 2j), where at further distances their speed began to converge with the control’s (Fig. 2k-l). This suggests that the synthetic cytokine receptors sensitize CAR-T cells to targeted factors secreted by neighboring cells, likely IL-5 in this case. Together, these data indicate the presence of highly motile subpopulations, induced by VRs, that sensitize CAR-T cells to self-secreted factors.

### VRs promote CAR-T cell motility through ROCK-myosin signaling, enabling migration through confined environments

Thus far, we have shown that VRs functionally push standard CAR-T cells into a high-motility phenotype in collagen I matrices. However, cells can navigate the ECM through multiple pathways, including ECM remodeling via metalloproteinases^22^, integrin-mediated focal adhesion formation^23^, or actomyosin-driven blebbing^24^. To determine the molecular mechanism resulting in this highly motile phenotype, we focused our investigation on the ROCK-myosin axis central to the amoeboid migration typically used by T cells^10,25,26^.

We began by staining the M5 and M5-VR5αIL5 cells for total myosin light chain (tMLC) and phosphorylated myosin light chain (pMLC) (Fig. 3a). CAR-T cells expressing the VR exhibited a 3-fold increase in tMLC and more than double the pMLC compared to standard CAR-T cells (Fig. 3b-c), supporting the mechanism whereby enhanced actomyosin contractility induces more blebbing and cellular deformation in the VR-CAR-T cells. To further support this, we created a PDMS-based microchannel device^27,28^ with widths varying from 3-50 microns and a fixed height of 10 µm and seeded M5 and M5-VR5αIL5 cells in the adjacent seeding areas to assess their ability to enter, or infiltrate, the microchannels (Fig. 3d). Given our proposed mechanism, inhibition of the ROCK-myosin pathway should impair the ability of the VR-CAR-T cells to enter the microfluidic channels (Fig. 3e). To test this, we treated both conditions with the ROCK inhibitor Y27632 during the microchannel migration assay. Inhibiting ROCK reduced VR-CAR-T cell entry into the channels by more than 4-fold compared to control levels, which remained unaffected by the inhibitor (Fig. 3f).

**Figure 3.**
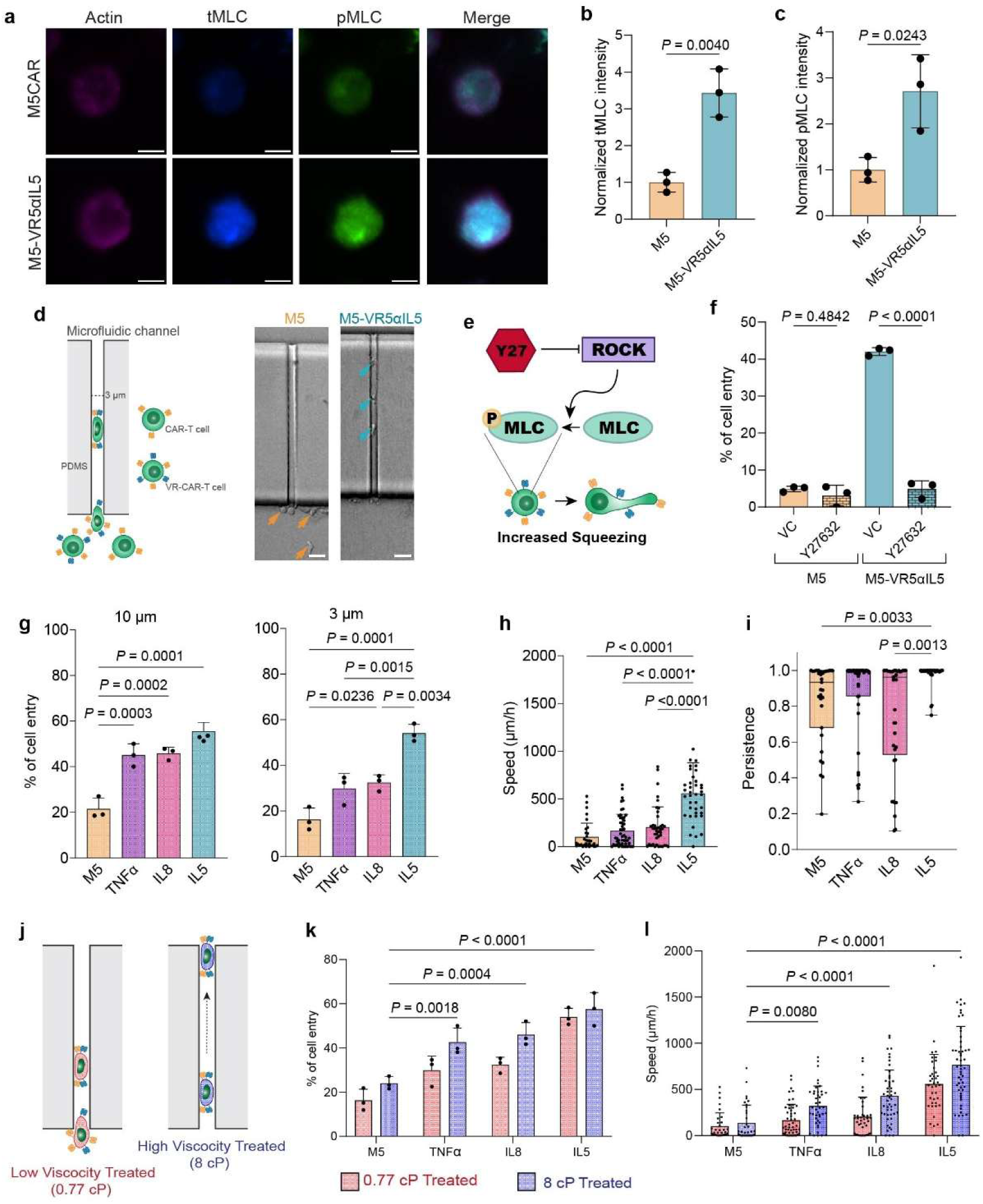
VR-mediated ROCK-myosin enrichment improves CAR-T cell confined motility across distinct cytokine-targeting receptors. (a) IF staining of CAR-T cells with and without a VR in 2D showing tMLC and pMLC expression. Scale bar = 5 µm. (b-c) Quantification of the IF staining showing pMLC and tMLC intensity normalized to M5. N = 3 independent experiments, n = 44 cells, Donors= 2. A Student’s t-test was used to assess statistical significance. (d) M5CAR and VR-CAR-T cells were seeded into microchannels and analyzed for percent entry and intrachannel motility parameters. Representative brightfield images show M5CAR and M5-VR5αIL5 CAR-T cells entering and migrating within microchannels (scale bar = 10 µm). (e) Schematic diagram of the ROCK-myosin pathway illustrating the target site of inhibition by Y-27632 (Y27). (f) VR-CAR and CAR-T cells were treated with the ROCK inhibitor Y27 and seeded into the microfluidic device. Y27 severely reduced the percentage of cell entry for VR-CAR-T cells while having no effect on the control. N = 3 independent experiments, Donors= 2. Statistical significance was assessed by one-way ANOVA with Sidak’s multiple comparisons test. (g) VR variants exhibit increased microchannel entry across multiple channel widths relative to the M5CAR control, consistent with enhanced invasive capacity. (h) Within channels, VR-expressing cells display increased migration speed, and (i) enhanced persistence compared to control cells. (g-i) N = 3 independent experiments. Statistical significance was assessed by one-way ANOVA with Tukey’s multiple comparisons test. (j–l) Treated under physiologically relevant viscosity conditions, VR-CAR-T cells demonstrate improved infiltration and confined migration relative to control. (k–l), N = 3 biological replicates. Statistical significance was assessed by two-way ANOVA with Tukey’s multiple comparisons test. (h, i, l) n > 30 cells per condition. (g-l) TNFα, IL8, and IL5 correspond to M5-VR5αTNFα, M5-VR5αIL8, M5-VR5αIL5, respectively. (b-l) All experiments performed at a width of 3 microns unless otherwise stated.

We next expanded our analysis to other variants of the VR, specifically, those that either target IL-8 or TNFα, in addition to IL-5, to evaluate whether this increase in confined motility is observed when targeting other autocrine secreted inflammatory factors. At widths of 10-50 microns, all VR-CAR-T cells had a higher percentage of cells that entered the microchannels compared to the M5 CAR control (no chemotactic gradient, Supplementary Fig. 3a). At a width of 3 microns, only cells expressing VRs targeting IL-8 and IL-5 had a higher percentage of cells that entered the channels (Fig. 3g). Beyond entry efficiency, cell motility within confined spaces is critical for rapidly infiltrating and eliminating disseminating tumor cells. At a confinement of 50 microns, both IL-8– and IL-5-dependent VR-CAR-T cells moved faster through the channels than M5CAR T cells. However, under tighter confinement (10 and 3µm), only IL-5-targeting VR-CAR-T cells maintained significantly greater speeds and velocities relative to controls (Figure 3h and Supplementary Fig. 3b-c).

In addition to speed, a cell’s capacity to migrate unidirectionally through the channel, or persistence, is essential for efficient extravasation and penetration. Accordingly, we quantified the directional persistence of VR-CAR-T cells under the same conditions. At 50 µm, both IL-8– and IL-5-targeting VR-CAR-T cells outperform the M5CAR control (Supplementary Fig. 3d). However, as with speed, at a 3 µm width only IL-5 outperformed the control (Fig. 3i).

Despite the role of confinement in regulating cell migration, prolonged exposure to other factors, such as extracellular fluid viscosity, also plays an active role in this process^27–29^. Compared to healthy controls, blood viscosity is significantly elevated in patients with advanced stages of cancer^30,31^ and both groups exhibit markedly higher viscosity than basal cell culture media. To account for this, we treated our CAR-T cells in a highly viscous medium of 8cP before seeding them into a 3-micron-wide channel to observe how these cells behave after treatment under more physiological conditions (Fig. 3j). At 3 microns, all VRs tested entered the channel more efficiently and moved faster under confinement than the M5CAR control, in contrast to previous conditions cultured in standard media (0.77cP), where only IL-8 and/or IL-5 demonstrated significant differences (Fig. 3k-l). Taken together, these data identify a novel mechanism by which cell infiltration is enhanced using synthetic cytokine receptors.

### CAR-T cells expressing VRs retain anti-tumor activity during TNBC exposure in vitro

To confirm that the VR-CAR-T cells retain their cytotoxicity against malignant breast cancer cells, we performed a cell killing assay using luciferase-tagged MDA-MB-231 and MCF7 cells, both of which express mesothelin^32,33^ (Supplementary Fig. 4a-c). All VR-CAR-T variants killed both cancer cell lines similarly to the standard M5CAR-T cells lacking the VR. Since all conditions were found to kill MDA-MB-231 cells at a higher efficiency than the MCF-7 cell line (Fig. 4a), all following experiments used the MDA-MB-231 cell line. ELISAs were then conducted on supernatants of the cell-killing assay to measure secreted TNFα and IFNγ, key cytokines expressed during T cell activation. Only IL-8 showed higher expression of the TNFα cytokine compared with the others. For IFNγ, secretion profiles were lower for all VR variants except for IL-8, which was comparable to the control (Fig. 4b). These data indicated that while IL-8-targeting VR-CAR-T cells exhibit robust pro-inflammatory responses in vitro, those targeting IL-5 and TNFα display comparatively attenuated inflammatory activity.

**Figure 4.**
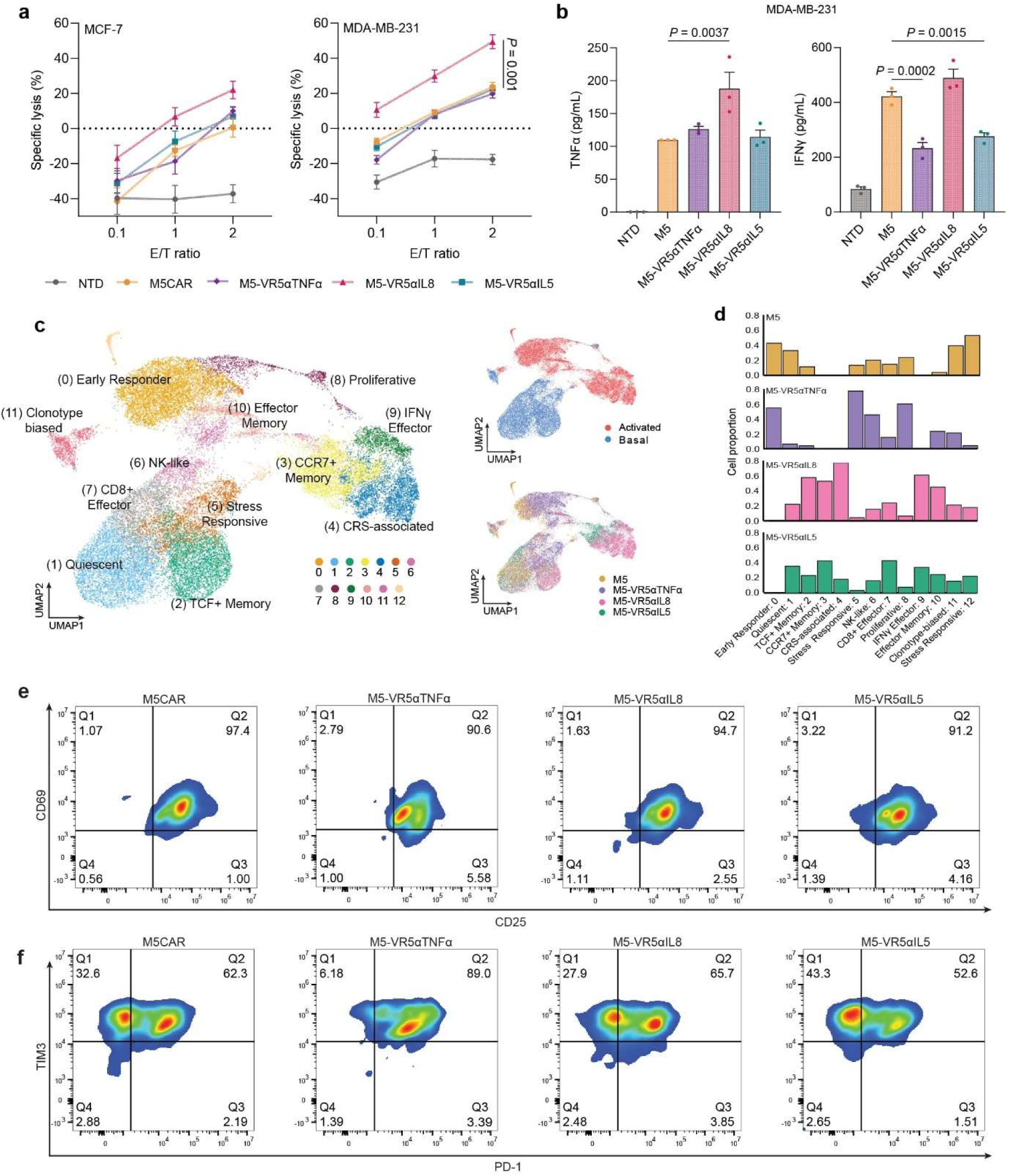
VRs enhance CAR-T motility without compromising antitumor function while exhibiting phenotypes associated with improved persistence compared to control CAR-T cells. (a) Cytotoxicity of M5 and M5-VR-T cells against MCF7 and MDA-MB-231 breast cancer cells 18 hours after coincubation at indicated effector/target (*E/T*) ratios. Data plotted as mean ± SEM, N=2 biological replicates, n=3 technical replicates. Two-way ANOVA with Tukey’s multiple comparison test was used to assess statistical significance. (*P<0.05, **P<0.01, ***P<0.001, P<0.0001). (b) ELISA assays measuring TNFα and IFNγ secretion by CAR and CAR-VR-T cells co-incubated with MDA-MB-231 cells at a 1:1 E:T ratio. Data plotted as mean ± SEM, N=2 biological replicates, n=3 technical replicates. One-way ANOVA with Tukey’s multiple comparison test was used for statistical significance. (*P<0.05, **P<0.01, ***P<0.001, P<0.0001). (c) Uniform Manifold Approximation and Projection (UMAP) visualization of high-quality single CAR-T cells with different CAR structures. Unsupervised clustering of both basal unstimulated and mesothelin-specific activated CAR-T cells identifies 13 distinct clusters, with cell type annotations indicated on the UMAP. (d) Frequency of each condition in a specific cluster of the UMAP space. (e,f) Flow cytometry analysis of CAR and CAR-VR cells co-incubated with MDA-MB-231 cells for 72 h. Frequency of cells expressing (c) CD25 and CD69 or (d) PD-1 and TIM3.

Using a publicly available scRNA-seq dataset and associated GitHub preprocessing pipeline of VR-CAR-T cells stimulated with mesothelin-expressing 3T3 cells or maintained under basal conditions^34^, we performed unsupervised clustering and visualized transcriptional heterogeneity using UMAP embeddings of jointly analyzed samples (Fig. 4c). Cells from M5, M5-VR5αIL5, M5-VR5αIL8, and M5-VR5αTNFa conditions occupied distinct but partially overlapping transcriptional programs in the shared UMAP space. Cluster annotation based on canonical marker genes and top differentially expressed transcripts revealed an enrichment of memory-associated transcriptional programs in VR-CAR-T cells targeting IL-5 or IL-8. Conversely, the M5 control contained the highest proportion of cells in early responders and stress-responsive clusters. Clusters upregulating genes for cytokines implicated in cytokine release syndrome (CRS) were predominantly of IL-8-targeting cells (Fig. 4d and Supplementary Fig. 4d). These data provide evidence for the enhanced persistence of both IL-5 and IL-8-targeting VRs. However, concurrent upregulation of genes associated with CRS, together with increased persistence, may promote accelerated exhaustion and increase the risk of adverse inflammatory effects in the IL-8-targeting cells.

To validate these insights regarding T-cell persistence, we co-cultured CAR-T cells with MDA-MB-231 cells and assessed activation and exhaustion profiles using CD69 and CD25 or TIM3 and PD-1 expression, respectively. Although activation was unchanged (Fig. 4e), the VR5αIL5 receptor induced a slight decrease in exhaustion compared to the control: VR5αIL5 cells had 52.6% exhausted cells, whereas the M5CAR had 62.3% (Fig. 4f). This suggests the potential for improved in vivo persistence relative to the other conditions. Together, this suggests a coordinated shift in activation, effector function, and persistence-associated phenotypes induced by VRs. These data suggest that VRs may promote transcriptional and functional states consistent with improved endurance during TME exposure.

### Enhanced CAR-T cells reduce primary and metastatic TNBC tumor burden

After validating the enhanced migration capabilities of the CAR-T cells, we next assessed how well they performed in a spontaneous metastatic model. Our hypothesis was that the enhanced confined migration of VR-CAR-T cells would allow for easier extravasation (Fig. 1e), where disassociated tumor cells could be eliminated at the primary tumor before disseminating to the lungs. MDA-MB-231 cells were injected orthotopically and allowed to establish for up to 27 days, after which the CAR-T therapies were administered intravenously via tail vein injection (Fig. 5a). Mice were monitored weekly to ensure healthy weights were maintained (Supplementary Fig. 5a). Mice of the tumor growth cohort were sacrificed 30 days after therapy administration to ensure tumor dissemination to the nearby organs, after which the lungs were harvested and assessed for metastatic burden using the novel CODA 3D mapping platform (Fig. 5b).

**Figure 5.**
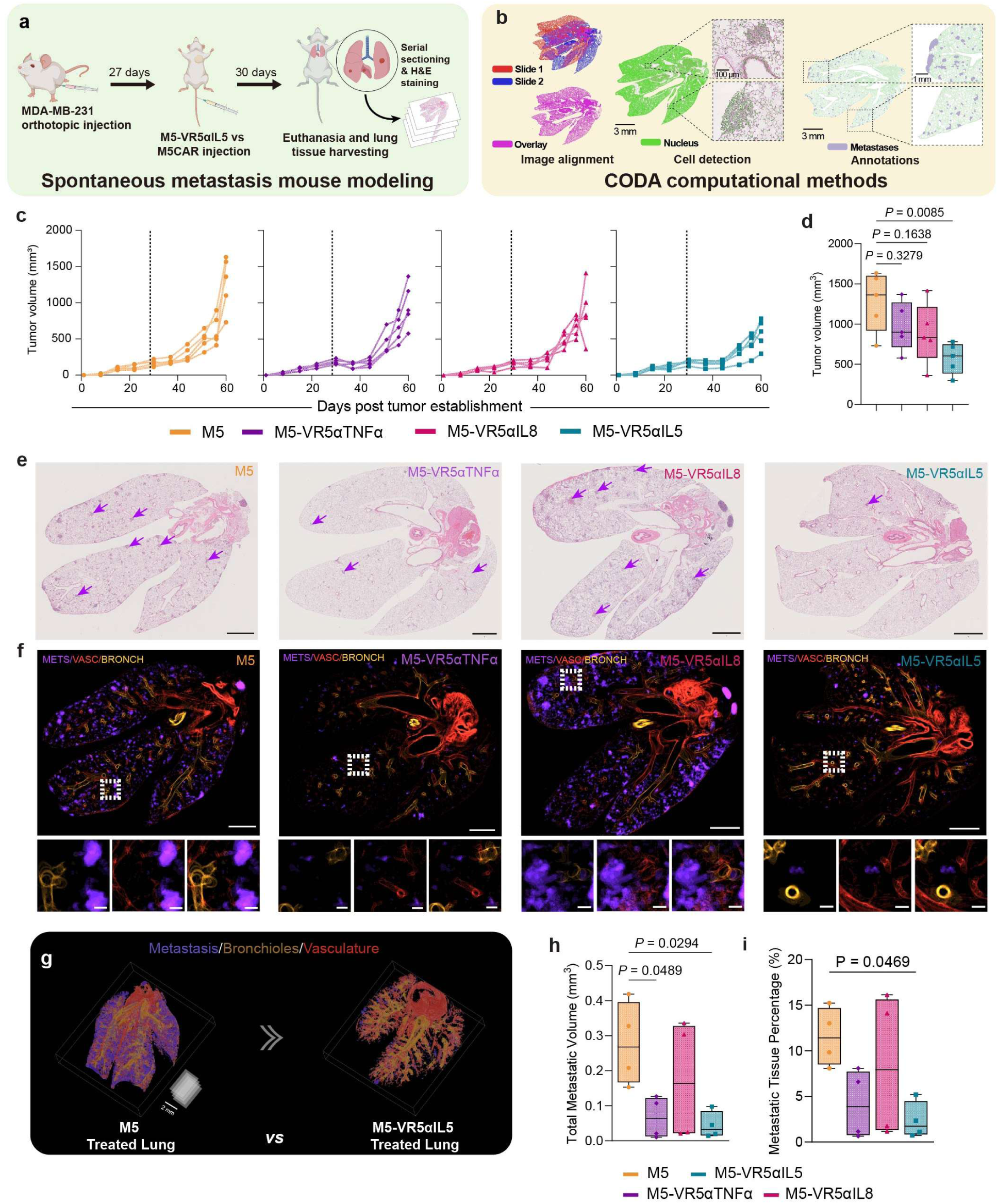
VR-CAR-T cells reduce metastatic tumor burden in an orthotopic CDX mouse model relative to M5CAR-T Cells. (a) NSG mice were injected orthotopically with MDA-MB-231 cells in the second left nipple of the mouse to establish a primary tumor. 27 days after injection, mice were treated with 3 million M5CAR-T cells expressing various velocity receptors (VRs). M5CAR-T cells that did not express VRs were the control. Mice were euthanized 30 days after CAR-T treatment. (b) After FFPE and H&E sectioning, consecutive sections were aligned before nuclei detection and structure annotations. (c, d) Administration of M5-VR5αIL5 CAR-T cells led to a significant reduction in the volume of the primary tumor compared to M5CAR-T cells. Other tested VRs resulted in minimal reductions compared to the control. n = 5 mice per group. One-way ANOVA with Tukey’s multiple comparison test was used for statistical significance. (*P<0.05, **P<0.01, ***P<0.001, P<0.0001). (e) Representative H&E scans of resected lung tissue from treated mice where purple arrows indicate metastatic lesions. Scale bar = 2mm. (f) Representative z-projections of resected lung tissue from treated mice. Regions in purple, red, and gold depict metastatic lesions, vasculature, and bronchioles, respectively. Scale bar = 2mm for whole tissue and 200 µm for zoomed-in structures. (g) Full 3D reconstruction consisting of 100 semi-consecutive slides of resected lungs from mice treated with M5CAR or the best-performing VR-CAR-T cell (h,i). Quantifications of total metastatic volume and metastatic tissue percentage. Data plotted as boxplots, n = 4 mice per group. One-way ANOVA with Tukey’s multiple comparison test was used for statistical significance. (*P<0.05, **P<0.01, ***P<0.001, P<0.0001).

Mice injected with the enhanced CAR-T therapy targeting IL-5 improved survival by reducing the volume of the primary tumor by 2-fold in mice afflicted with the disease (Fig. 5c-d and Supplementary Fig. 5b-d). Due to the extensive time period between therapy injections and organ harvesting, no CAR-T cells were found in IHC-stained tumor sections (Supplementary Fig. 5e).

The CODA platform developed is based on a deep learning model trained to identify metastasis (purple), bronchioles (yellow), and vasculature (red) based on H&E images of the lungs (Fig. 5e-f and Supplementary Fig. 6a-d). The deep learning model was applied to stacks of 15 consecutive slides for each condition to produce maximum intensity Z projections (Fig. 5f). The analysis revealed that the lungs in M5CAR-treated mice contained the largest number and volume of metastatic lesions, which constituted a larger percentage of the lung tissue compared to mice treated with VR-CAR-T therapies. The M5-VR5αIL5 condition performed the best, lowering the median total metastatic volume by more than 8-fold and the median metastatic tissue percentage by more than 6-fold compared with the control (Fig. 5g-i). When correlating primary tumor volume, average metastatic lesion size, and the number of metastatic lesions, the M5-VR5αIL5 condition consistently showed the lowest values across all three parameters (Supplementary Fig. 6e-f). In sum, supplementing standard CAR-T cells with VRs targeting IL5 shows a remarkable effect not only against the solid tumor, but also against distant organ metastasis.

To further resolve transcriptional differences between control and the top-performing VR-CAR-T cells that contribute to the observed in vivo outcomes, we reanalyzed the single-cell RNA-seq dataset, restricting the analyses to the M5 and M5-VR5αIL5 conditions. Unsupervised clustering, as performed previously, and cluster identities were assigned based on canonical marker genes and the top five differentially expressed genes per cluster, visualized using a heatmap. (Supplementary Fig. 7a-c). Basal populations were primarily composed of quiescent and NKG2D⁺ effector cells, whereas activated populations included early responder, stress-responsive, proliferative, IFNγ effector, and activated memory-like subsets. Notably, the M5-VR5αIL5 condition contributed disproportionately to several key clusters, including activated memory-like, NKG2D⁺ cytotoxic effector, and quiescent populations, suggesting a more functionally diverse cellular landscape. To better understand intercellular communication, we performed a CellChat analysis and visualized signaling interactions using chord plots (Supplementary Fig. 7d). These results demonstrated strong connectivity between activated memory-like, NK-like cytotoxic, and NKG2D⁺ effector populations, indicating active crosstalk among cytotoxic and functionally engaged subsets. In addition, quiescent cells exhibited extensive interactions with multiple clusters, including proliferative, activated memory-like, and early responder populations, suggesting their role as a central signaling hub within the network.

Finally, to specifically interrogate migratory potential, we curated 20 migration-related gene sets from GSEA (Supplementary Table 1) and performed unsupervised clustering using only genes from these datasets. Unsupervised clustering was performed using the Leiden algorithm following graph-based construction of a shared nearest-neighbor network. This produced a UMAP embedding in which clusters were annotated based on the highest gene set enrichment ratios (Supplementary Fig. 7e). Five gene sets showed the most distinct clustering patterns: vascular endothelial cell (VEC) adhesion, integrin signaling, extravasation, bleb assembly, and leukocyte migration. Within this framework, the M5-VR5αIL5 condition exhibited the highest proportion of cells within extravasation and leukocyte migration clusters (Supplementary Fig. 7f-g). Taken together, these results suggest that M5-VR5αIL5 CAR-T cells are not only transcriptionally primed for migration but also inflammatory programs that contribute to their anti-tumor response.

## Discussion

Chimeric antigen receptor (CAR) T cell therapy has shown limited efficacy in solid tumors, where mortality is driven largely by late-stage dissemination of cancer cells and the formation of metastases. A critical yet underemphasized requirement for effective solid-tumor CAR-T therapy is the ability to prevent and control metastatic spread in addition to reducing primary tumor burden. Here, we demonstrate that engineering CAR-T cells with a synthetic cytokine receptor that enhances migratory capacity (VR) improves both intratumoral function and antimetastatic activity in a highly metastatic TNBC model.

Using the CODA platform, we observed that VR-expressing CAR-T cells significantly decreased lung metastases compared with CAR-T cells lacking the receptor, in correlation with a reduction in size of the primary tumor. These in vivo outcomes were consistent with the superior migratory performance of VR-CAR-T cells under physical constraints. In confined microchannels of varying widths, VR-CAR-T cells migrated more efficiently relative to controls. In 3D collagen matrices, we found that VR-CAR-T cell motility depended on local cell proximity, with key trajectory features such as higher speed, MSD, and net displacement enriched in VR-dominant clusters. Single-cell transcriptional profiling of migration-relevant gene expression datasets further showed that VR-CAR-T cells were most enriched for programs related to extravasation and leukocyte migration. Notably, similar immunological profiling identified memory-associated clusters among the most effective VR-CAR-T conditions (e.g., M5-VR5αIL5), suggesting that both enhanced motility and favorable differentiation states contribute to efficacy.

Given that most CAR-T trials for solid tumors enroll patients with advanced or relapsed disease, and therefore a high likelihood of metastatic involvement^35^, therapies must be capable of controlling both primary tumors and disseminated lesions. While FDA approvals currently remain limited to hematologic malignancies, our results provide proof of concept that engineering cell-intrinsic motility and extravasation can improve trafficking to tumor sites and reduce metastatic burden. In our study, the best-performing VR condition reduced total metastatic volume and incidence by more than 4-fold, likely through a combination of improved infiltration of the primary tumor and early elimination of locally disseminating cancer cells before intravasation.

The tumor microenvironment presents multiple barriers to CAR-T function, including physical obstacles (e.g., basement membrane)^8,36^, suppressive cytokines (e.g., TGF-β, IL-6)^4^, and inhibitory checkpoints (e.g., PD-L1, TIGIT)^37^ that collectively drive exhaustion and limit persistence. Among VR designs tested (e.g., TNFα-, IL-8-, or IL-5–responsive constructs), the top performer was the one that combined superior infiltration under confinement with enhanced 3D migratory capacity. This condition aligned with transcriptional clusters enriched for memory signatures, consistent with prior reports linking memory-associated phenotypes with durable responses in some patients^38–40^. Together, these data support a model in which motility, extravasation, and persistence are jointly associated with stronger anti-tumor activity.

We acknowledge that additional determinants of efficacy in metastatic solid tumors, such as resistance to immunosuppression^4,12,37^, optimal antigen density recognition^41^, and metabolic fitness^42^, exist but were not the primary focus of this work. Nonetheless, our findings highlight motility engineering via synthetic cytokine receptors as a compelling strategy to enhance both tumor infiltration and metastatic control. Future studies should prospectively test whether VR-driven migratory phenotypes correlate with improved clinical outcomes and evaluate combination strategies (e.g., checkpoint blockade, matrix remodeling, or metabolic support) to further potentiate anti-tumor efficacy.

## Acknowledgments

The authors thank Jyothsna Girish for her invaluable support and feedback throughout this work. The authors would also like to thank prior and current members of the Wirtz lab for their guidance and support. The authors acknowledge support from the Integrated Imaging Center and Oncology Tissue Services at Johns Hopkins University. All figures were made using BioRender and Adobe Illustrator.

## Materials and Methods

### Cell lines

Human breast cancer MCF-7 (HTB-22, ATCC) and triple-negative breast cancer MDA-MB-231 (CRM-HTB-26, ATCC) served as representative breast cancer cell models to examine the effects of VR-CAR-T therapy. HEK293T cells (CRL-1573, ATCC) were used in the production of lentiviruses. All human cell lines were maintained in DMEM supplemented with 10% FBS and 1% penicillin/streptomycin. MDA-MB-231 cells were engineered with lentiviruses to express luciferase-P2a-mCherry and sorted using a Sony SH800 cell sorter to isolate mCherry-positive cells.

### Lentivirus production

CAR and VR constructs were cloned into the backbone of the vector pSLCAR-CD19-BBz (Addgene) using Takara’s in-fusion assembly. CAR receptors were cloned upstream of the P2A-GFP sequence and downstream of the EF1a promoter after removal of the original backbone’s CD19 CAR sequence. VR receptors were clones downstream of the EF1a promoter after removal of original backbone’s CD19 CAR-P2A-GFP sequence. Lentiviral particles were produced by transfecting HEK293 T cells with plasmids containing either the M5CAR sequence or the VR sequence along with plasmids PSPAX and pMD2.G using GeneJuice (Millipore Sigma) in Opti-MEM supplemented with 5% FBS. Particles were isolated through centrifugation using Lenti-X Concentrator.

### CAR-T cell generation

Pan-human T cells were isolated from PBMCs using the STEMCELL Technologies negative selection kits before overnight activation using CD3/CD28 Dynabeads in X-Vivo supplemented with 5% Human AB serum, 1% P/S (Complete X-Vivo) and 100 IU/mL of IL2. The following day, T cells were transduced with lentiviruses for the VRs for 2 days on Retronectin-coated plates at 1E6 cells/mL. Transduction efficiency of the VRs was then checked by staining 50uL of cells with a F(ab)2 antibody (Jackson Immunoresearch), and the remaining cells were transduced with the M5 CAR virus for another 2 days. Cells were then washed twice with HBSS before being allowed to rest in complete X vivo supplemented with 100 IU of IL2 before Dynabead removal at days 5-7. Following this period, eGFP expression was quantified by flow cytometry as a proxy for CAR expression before downstream analyses.

### Cytotoxicity assays

Luciferase-tagged cancer cells were seeded in white walled 96 well plates at 4000 cells per well and incubated overnight at 37°C and 5% CO2. The next day, CAR-TV cells were added to each well at the indicated E:T ratios, and allowed to react overnight. The next day, 100 μL of supernatant was removed from each well, and 100 μL of Bright glow substrate was added and shaken for 5 min at 400 RPM. Seven minutes later, relative light units were measured on a SpectraMax plate reader (Molecular Devices) at an integration time of 500 and 1000 ms.

### Activation/exhaustion phenotyping

VR-CAR-T cells were co-incubated with MDA-MB-231 cells at a 2:1 E:T ratio for 3 days at 37°C and 5% CO2. Culture supernatant containing T cells was then removed from each well before being washed with FACS buffer. Cells were then stained with 1:200 of the following antibodies: anti-CD25 BV510, anti-CD69 BV421, anti-PD-1 PE, and anti-TIM3 BB700 (for 20 minutes at 4°C in the dark. Cells were washed 3X times before being analyzed on a Cytoflex and gated on eGFP to assess activation and exhaustion of CAR-positive cells.

### Cytokine assays

To further measure T cell activation, MDA-MD-231 cells were seeded at 4000 cells per well of a 96-well plate with T cells added on top the following day at a 1:1 E:T ratio. After an overnight incubation, 100 μL of the culture supernatant was removed and analyzed by an IFNγ and TNFα ELISA (Invitrogen) using the manufacturer-specified protocol.

### 3D migration assays

Cell-embedded 3D collagen I gels were prepared according to a previously described method^19^. After the 5-7 day activation period, CAR-T cells were resuspended in complete X-Vivo and embedded in a 2 mg/mL collagen I mixture consisting of HBSS, rat tail collagen 1, 0.2 M NaOH, and 100 IU/mL of IL2. The cell suspension mixture was then plated at 100uL/well in a 96-well plate and allowed to solidify at 37°C and 5% CO2 for 1.5 hours. X-Vivo supplemented with 100 IU/mL of IL2 was then added on top of each well before incubating for 24 hours. Following this period, gels were imaged every 2 min for 1 h using a Nikon Ti2 microscope equipped with a live-cell system to maintain the gels at 37°C and 5% CO_2_. Time-lapsed images were viewed on Fiji Image J and cells tracked using MTrackJ to determine cell trajectories, as well as basic migration-associated parameters.

### Single-cell transcriptome data processing and analysis

Publicly available single-cell RNA-seq datasets of mesothelin-stimulated and basal CAR-T cells were obtained from a publicly available GitHub repository (Velocity-CAR-T) developed by Zhiliang Bai along with the associated preprocessing pipeline. Processed Seurat objects corresponding to stimulated (MESO) and basal (Basal) conditions were imported as RDS files and merged into a single Seurat object for downstream analysis. Default RNA assays were retained for initial integration and metadata harmonization. CAR-T cell subsets corresponding to selected receptor constructs were isolated based on annotated structure labels (e.g., M5, IL-5). Downstream analyses were performed on this subset. Cells were normalized and variance-stabilized using SCTransform, with percent mitochondrial gene expression regressed out.

Principal component analysis (PCA) was performed on the SCTransform-normalized data, and the top principal components (1–25) were selected based on elbow plot inspection. A shared nearest neighbor graph was constructed using these PCs, followed by unsupervised clustering using the Louvain algorithm at a resolution of 0.4. Uniform Manifold Approximation and Projection (UMAP) was used for two-dimensional visualization of transcriptional states. Cellular composition across clusters and experimental conditions was quantified by calculating the proportion of cells from each CAR-T cell condition within each cluster. These distributions were visualized as stacked bar plots to assess enrichment patterns across transcriptional states.

Cluster-specific marker genes were identified using Seurat’s FindAllMarkers function, retaining genes with a minimum of 10% expression in at least one group and a log fold-change threshold of 0.5. The top five differentially expressed genes per cluster were selected based on average log fold-change.

### Trajectory preprocessing

To ensure consistency across samples and reduce variability due to differences in trajectory length and orientation, all cell trajectories were normalized to a fixed duration of 1 h. Trajectories with missing time points were discarded, and only cells continuously tracked throughout the entire interval were included in downstream analyses to maintain data reliability. To address directional variability, we defined an intrinsic migration coordinate system. The principal migration axis was determined as the direction of maximal cumulative displacement, while the secondary axis was defined orthogonally^43,44^. These axes were obtained via singular value decomposition (SVD) of the displacement matrix *A*, such that:

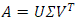

where *U* and *V*^T^ represent the eigenvector matrices of *AA*^T^ and *A*^T^*A*, respectively, and Σ contains the singular values. The original coordinate matrix *R* was then projected onto this orthonormal basis via *V*, such that:

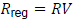

In this coordinate system, the first and second dimensions correspond to motion along the primary and secondary migration axes, respectively.

### Multidimensional motility analysis

To systematically quantify single-cell motility behavior, we extracted a set of 88 features from 262 cells, capturing multiple aspects of movement dynamics^27,45^. These features included both displacement-based and turning angle–based metrics, encompassing characteristics such as movement magnitude, spatial and temporal organization, signal properties, correlation structure, and decomposed motion components. A full description of these features is provided in Supplementary Table 2. To reduce redundancy among correlated variables, principal component analysis (PCA) was performed, and components explaining 95% of the total variance were retained. These principal components were subsequently used for two-dimensional visualization via UMAP, followed by unsupervised clustering. Clustering was conducted using the K-means++ algorithm, and cluster quality was evaluated by examining the projected trajectories associated with each motility cluster for qualitative coherence. Clusters containing fewer than 10 cells were excluded to improve robustness, resulting in four final motility clusters.

### Migration in microfluidic devices

Microfluidic devices were fabricated as previously described^24,46,47^. Briefly, silicon wafers were patterned by photolithography to generate arrays of parallel microchannels. Each channel measured 200 µm in length and 10 µm in height, with widths of 3, 6, 10, 20, or 50 µm. Polydimethylsiloxane (PDMS; Ellsworth Adhesives, #2065622) was poured over the patterned wafers, degassed under vacuum for 30 min to remove trapped air, and then cured at 85°C for ∼90 min. Once polymerized, six wells for cell seeding and media exchange were created using a 5 mm biopsy punch. The PDMS layers were immersed in 100% ethanol and sonicated for 5 min, followed by thorough rinsing in deionized water. PDMS slabs and glass slides (25×75 mm; Electron Microscopy Sciences, #72192-75) were each plasma-treated (PDC-32G, Harrick) to activate their surfaces and then carefully aligned to achieve irreversible bonding. After assembly, devices were coated with rat tail collagen type I (20 µg/mL; Gibco, #A1048301) and incubated at 37°C for 1 h before use.

After bonding, approximately 50,000 cells (1,000 cells/µL) were loaded into the inlet of the seeding channel, allowing a pressure-driven flow to carry the suspension across the seeding region. When the advancing front reached the outlet, 6.67 µL of the suspension were withdrawn from the outlet port to halt further flow. Devices were then placed in a 37°C, 5% CO₂, fully humidified incubator for ∼30 min to allow cells to attach to the collagen I-coated seeding area adjacent to the microchannel entrances. Following attachment, fresh medium was added uniformly across all six wells, and migration experiments were initiated.

Time-lapse imaging was performed at 2-min intervals for 3–5 h using an inverted Nikon Eclipse Ti microscope (Nikon, Tokyo, Japan) equipped with motorized stage control and NIS-Elements acquisition software. A 10×/0.45 Ph1 objective was used for all recordings. Environmental conditions were maintained at 37°C and 5% CO₂ with a stage-top incubator (Okolab or Tokai Hit). Cell trajectories were obtained by manual tracking in ImageJ (NIH, Bethesda, MD) using the MTrackJ plugin. Subsequent analyses, including calculations of average velocity, instantaneous directional persistence, were performed in MATLAB (R2022b; MathWorks, Natick, MA).

### Immunofluorescence staining

Immunostaining was performed as previously described^48^. Briefly, cells were fixed with 4% paraformaldehyde (J19943K2, Thermo Fisher Scientific) for 15 min, washed three times with 1× PBS, and permeabilized with 1% Triton X-100 (Millipore Sigma) dissolved in 1× PBS for 10 min. Cells were then blocked for 1 h with 2% BSA and 0.1% Triton X-100 and incubated overnight at 4°C with primary antibodies against phosphorylated myosin light chain (pMLC; 1:200; 3671S) or total myosin light chain (tMLC; 1:200; AB89594), diluted in blocking buffer. Cells were washed three times with 1× PBS and incubated for 1 h at room temperature with the appropriate fluorescently conjugated secondary antibodies and Alexa Fluor 647 phalloidin (1:1000; A22287, Thermo Fisher Scientific) diluted in 2% BSA, 0.1% Triton X-100, and 2% goat serum. Finally, cells were washed three times for with 1× PBS and imaged using a Nikon AXR confocal microscope equipped with a Plan Apo 40×/1.15 WI (AXR) objective, respectively. Images were analyzed using ImageJ.

### In vivo mouse modeling

All animal experiments were approved by the institutional Animal Care and Use Committee (IACUC) of Johns Hopkins University (Protocol MOE23E29). Mice were maintained at no more than 5 mice per cage for all groups. Studies were performed in 8-12 week-old NOD-*scid* IL2Rgamma^null^ (NSG) female mice purchased from Johns Hopkins Medical Institution and housed under a 12-hour dark/light cycle at 25°C. MDA-MD-231 (1×10^6^) human triple-negative breast cancer cells were suspended at a 1:1 ratio of Matrigel and DPBS and orthotopically injected into the left mammary fat pad to form breast tumors. Once tumors became palpable at a volume of around 100-250 mm³, CAR-T or VR-CAR-T (3×10^6^) cells suspended in 100 μL of DPBS were injected into the tail vein of the mouse using a 30-gauge syringe. Tumors were measured twice a week using digital calipers, and volume calculated using the following formula (*L x W*^2^) × 0.5. Half of the cohort was used for tumor growth studies, and the other used for a survival study, where each mouse was euthanized once the tumor volume reached 1500 mm³. Tumor and lungs were collected at the indicated time points, before being embedded in paraffin for H&E staining.

### Sample acquisition, serial sectioning and staining

Samples were processed into FFPE tissue blocks at the Johns Hopkins Oncology Tissue Core. Every other section was stained, resulting in approximately 15 H&E-stained slides and 15 unstained sections for each mouse tissue block. Unstained sections were stored in the –20°C under optimal humidity conditions. H&E slides were scanned using a Hamamatsu Nanozoomer S210 and saved as NDPI files. Files were converted to one, two, and four micron per pixel resolution TIFF files, using Openslide software^49,50^.

### Deep learning segmentation of mouse lung microanatomy

We annotated the mouse lung microanatomy by training two deep learning models and subsequently integrating their outputs^51^. The first model labelled components in mouse anatomy such as vasculature, bronchiole, alveoli, and extracellular matrix in all whole slide images of each mouse tissue block. The second model was trained to identify metastasis in the mouse lungs and nondiseased tissue. Models were combined to fully annotated the mouse lungs, allowing the integration and analysis of metastasis with the remaining anatomical components in the lungs.

### 3D mapping of metastasis and tissue components in mouse lungs

To better understand the effects of CAR-T therapies in mouse lungs, we produced a three-dimensional assessment of all microanatomical components^52^. H&E images were aligned using CODA methods, beginning with a global alignment to translate tissue sections, followed by a local elastic alignment that refined positioning to achieve micron-level accuracy^49^. InterpolAI was used to generate intervening missing H&E images and further enhance the microanatomical connectivity in 3D^53^

### Software

All migration analyses were performed in Python with the following software specifications: python 3.9.13, scipy 1.13.1, scikit-learn 1.1.13, scikit-image 0.19.3, scikit-posthocs 0.9.0, statsmodels 0.13.5, pandas 1.5.2, matplotlib 3.6.2, seaborn 0.11.2, umap-learn 0.5.3, and numpy 1.23.5, cmcrameri 1.9, EntropyHub 0.2, pyarrow 12.0.1.

## Statistical analysis

All statistical tests were performed using GraphPad Prism. All analyses were performed using either the Student’s t-test for two groups or the ordinary one-way ANOVA with Tukey’s post-test for comparisons of multiple groups.

## Data Sources

“Single-cell transcriptomic data of velocity-engineered CAR-T cells” were obtained from a publicly available GitHub repository (Velocity-CAR-T) developed by Zhiliang Bai^34^. The dataset includes single-cell RNA sequencing profiles of CAR-T cells expressing synthetic ‘velocity receptors’ designed to enhance migratory behavior. Data were downloaded from https://github.com/Zhiliang-Bai/Velocity-CAR-T and processed using custom Python scripts.”

