## Supplementary Materials for "Reprogramming CAR-T cell function through synthetic cytokine receptors suppresses metastatic progression of TNBC"

This file includes:

Figures S1 to S7 for multiple supplementary figures.

Tables S1 to S2 for multiple supplementary tables.

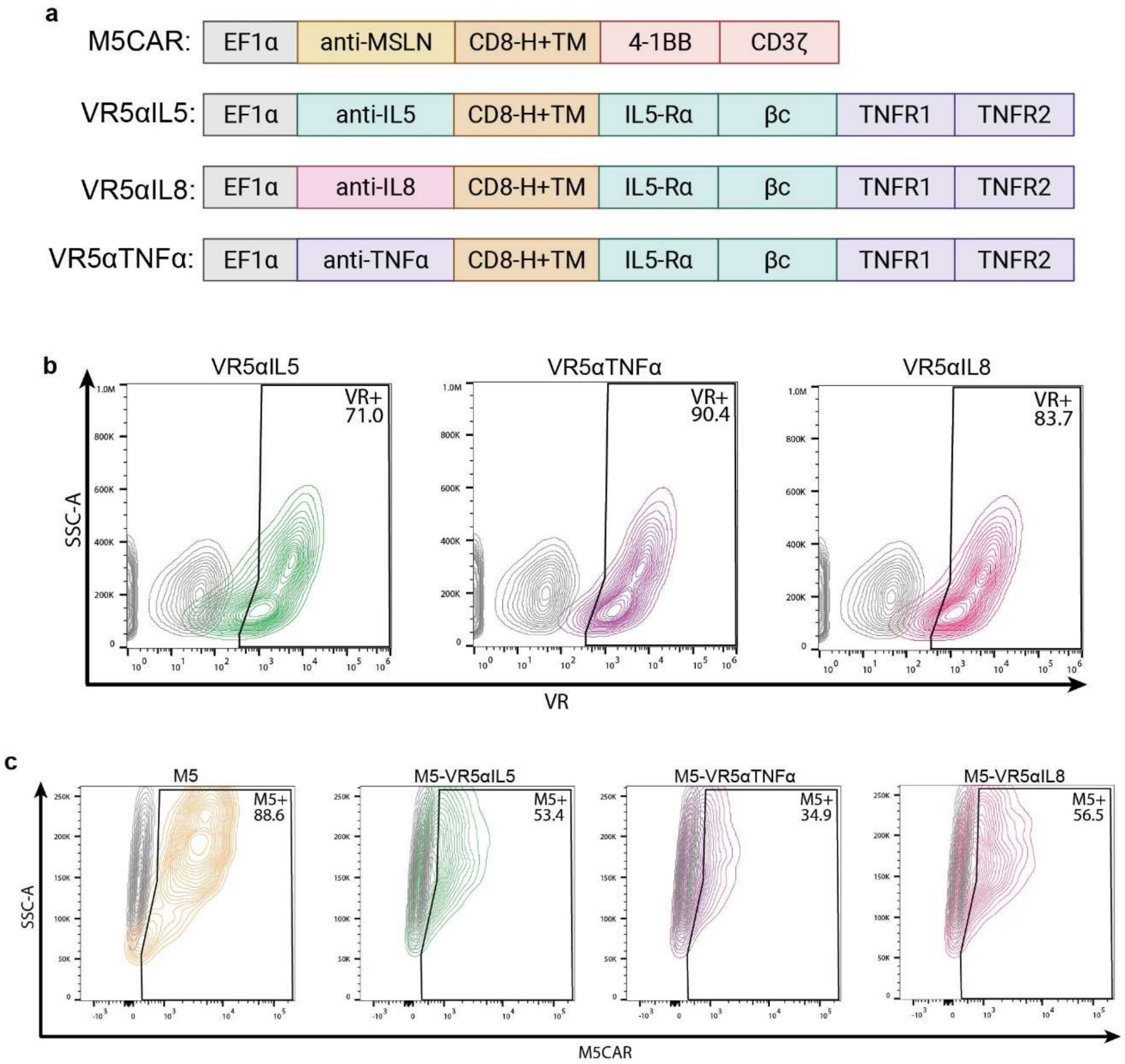


**Supplementary Figure 1. VR-CAR-T design and expression.** (a) Schematic illustration of M5CAR and synthetic VR constructs. (b) Expression of all synthetic VRs in T cells. T cells were transduced after overnight activation with CD3/CD28 Dynabeads. (c) Expression of M5CAR in previously VR-transduced T cells. All cells were co-incubated with the virus for 2 days during the second transduction.

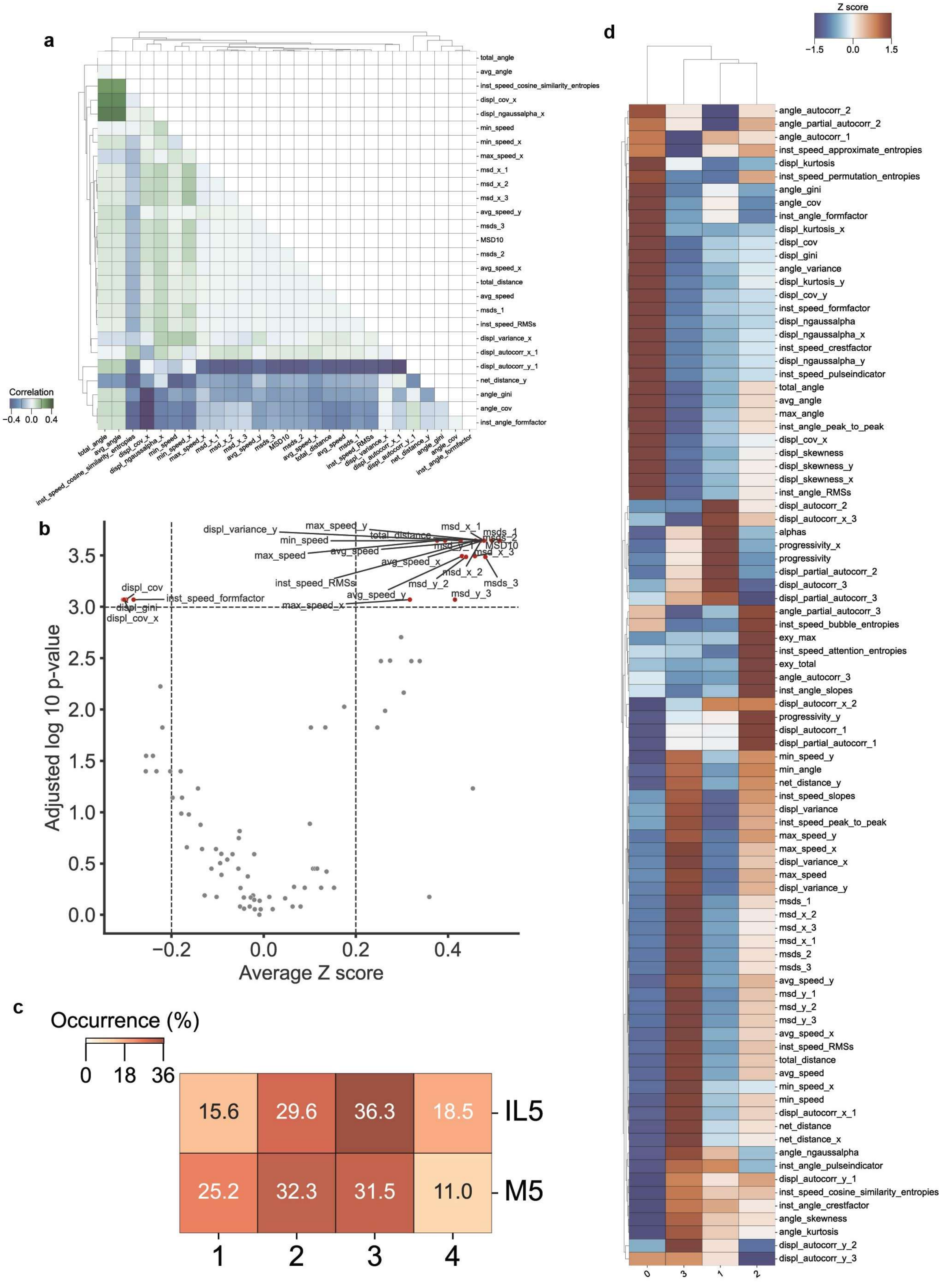
a
total_angle
avg_angle
inst_speed_cosine_similarity_entropies
displ_cov_x
displ_ngaussalpha_x
min_speed
min_speed_x
max_speed_x
msd_x_1
msd_x_2
msd_x_3
avg_speed_y
msds_3
MSD10
msds_2
avg_speed_x
total_distance
avg_speed
msds_1
inst_speed_RMSs
displ_variance_x
displ_autocorr_x_1
displ_autocorr_y_1
net_distance_y
angle_gini
angle_cov
inst_angle_formfactor
Correlation
−0.4 0.0 0.4
b
Adjusted log 10 p-value
Average Z score
displ_variance_y
max_speed_y
min_speed
max_speed
total_distance
avg_speed
avg_speed_x
inst_speed_RMSs
msd_x_1
msds_1
msds_2
MSD10
msd_y_1
msd_x_3
msd_x_2
msd_y_2
msds_3
msd_y_3
avg_speed_y
max_speed_x
displ_cov
inst_speed_formfactor
displ_gini
displ_cov_x
c
Occurrence (%)
0 18 36
15.6 29.6 36.3 18.5 IL5
25.2 32.3 31.5 11.0 M5
1 2 3 4
d
Z score
−1.5 0.0 1.5
angle_autocorr_2
angle_partial_autocorr_2
angle_autocorr_1
inst_speed_approximate_entropies
displ_kurtosis
inst_speed_permutation_entropies
angle_gini
angle_cov
inst_angle_formfactor
displ_kurtosis_x
displ_cov
displ_gini
angle_variance
displ_kurtosis_y
displ_cov_y
inst_speed_formfactor
displ_ngaussalpha
displ_ngaussalpha_x
inst_speed_crestfactor
displ_ngaussalpha_y
inst_speed_pulseindicator
total_angle
avg_angle
max_angle
inst_angle_peak_to_peak
displ_cov_x
displ_skewness
displ_skewness_y
displ_skewness_x
inst_angle_RMSs
displ_autocorr_2
displ_autocorr_x_3
alphas
progressivity_x
progressivity
displ_partial_autocorr_2
displ_autocorr_3
displ_partial_autocorr_3
angle_partial_autocorr_3
inst_speed_bubble_entropies
exy_max
inst_speed_attention_entropies
exy_total
angle_autocorr_3
inst_angle_slopes
displ_autocorr_x_2
progressivity_y
displ_autocorr_1
displ_partial_autocorr_1
min_speed_y
min_angle
net_distance_y
inst_speed_slopes
displ_variance
inst_speed_peak_to_peak
max_speed_y
max_speed_x
displ_variance_x
max_speed
displ_variance_y
msds_1
msd_x_2
msd_x_3
msd_x_1
msds_2
msds_3
avg_speed_y
msd_y_1
msd_y_2
msd_y_3
avg_speed_x
inst_speed_RMSs
total_distance
avg_speed
min_speed_x
min_speed
displ_autocorr_x_1
net_distance
net_distance_x
angle_ngaussalpha
inst_angle_pulseindicator
displ_autocorr_y_1
inst_speed_cosine_similarity_entropies
inst_angle_crestfactor
angle_skewness
angle_kurtosis
displ_autocorr_y_2
displ_autocorr_y_3
0 3 1 2

**Supplementary Figure 2. Features of VR-CAR-T motility.** (a) Matrix of correlation differences between M5-VR5αIL5 and M5. Green shades represent stronger correlation towards M5-VR5αIL5, and blue shades represent stronger correlation towards M5. Only features where correlation > |0.3| are shown. (b) Volcano plot of differentially regulated motility features between conditions. (c) Heat chart showing the proportion of all clusters in each condition. (d) Heat map showing Z-score expression of different motility features in each cluster of the UMAP space. Positive Z-scores indicate upregulation in that specific motility feature compared to the other clusters. (a-d) N = 3 biological replicates, M5: n = 127, VR5αIL5-M5: n = 135 cells.

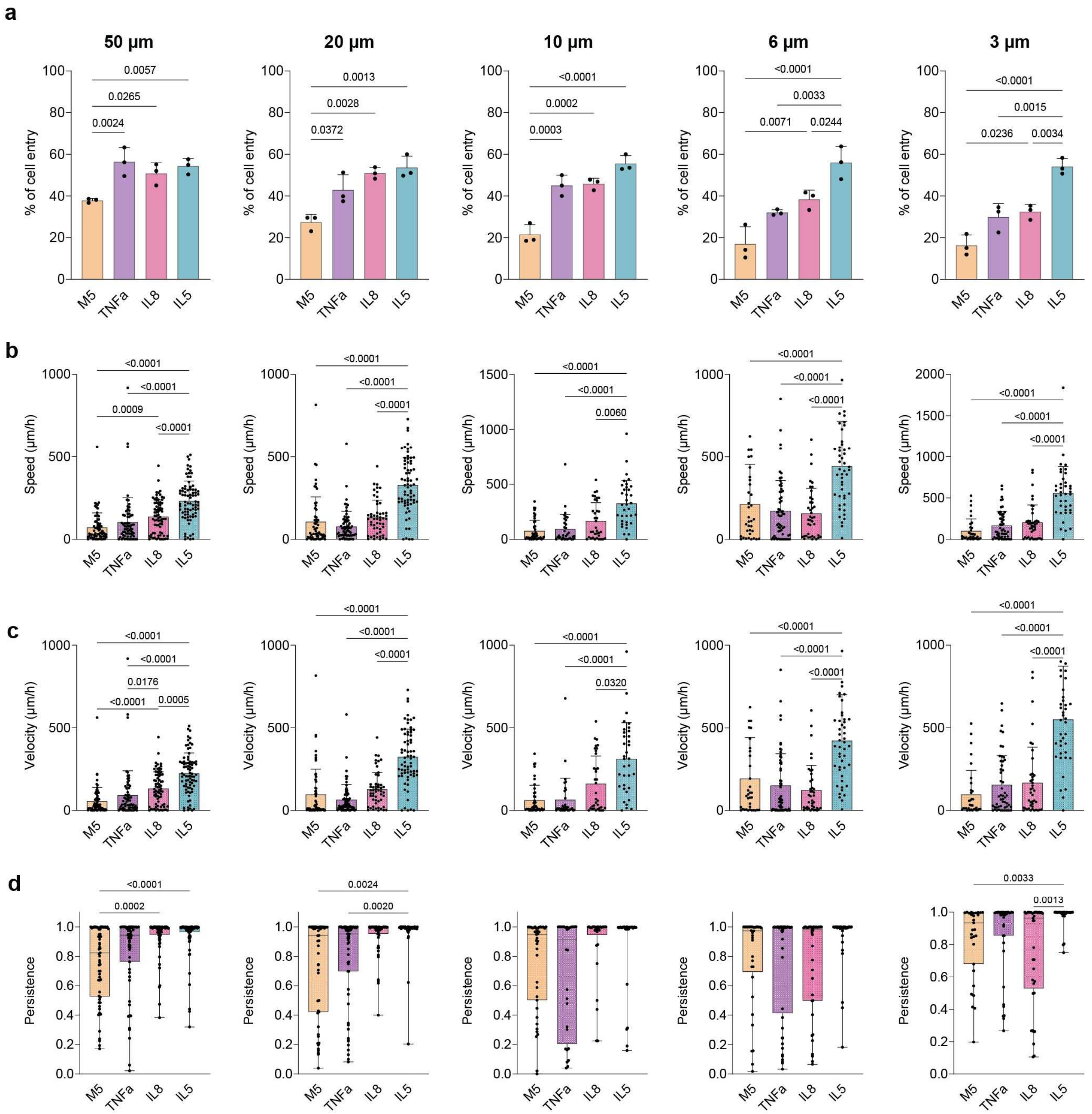


**Supplementary Figure 3. Motility within microchannels.** (a) VR-CAR T cells exhibit increased microchannel entry across multiple channel widths relative to the M5CAR control, consistent with enhanced invasive capacity. (b) Within channels, VR-expressing cells display increased migration speed (c) velocity and (d) enhanced persistence compared to control cells across multiple channel widths. (a-d) N = 3 biological replicates. Statistical significance was assessed by one-way ANOVA with Tukey's multiple comparisons test (*P < 0.05, **P < 0.01, ***P < 0.001, ****P < 0.0001).

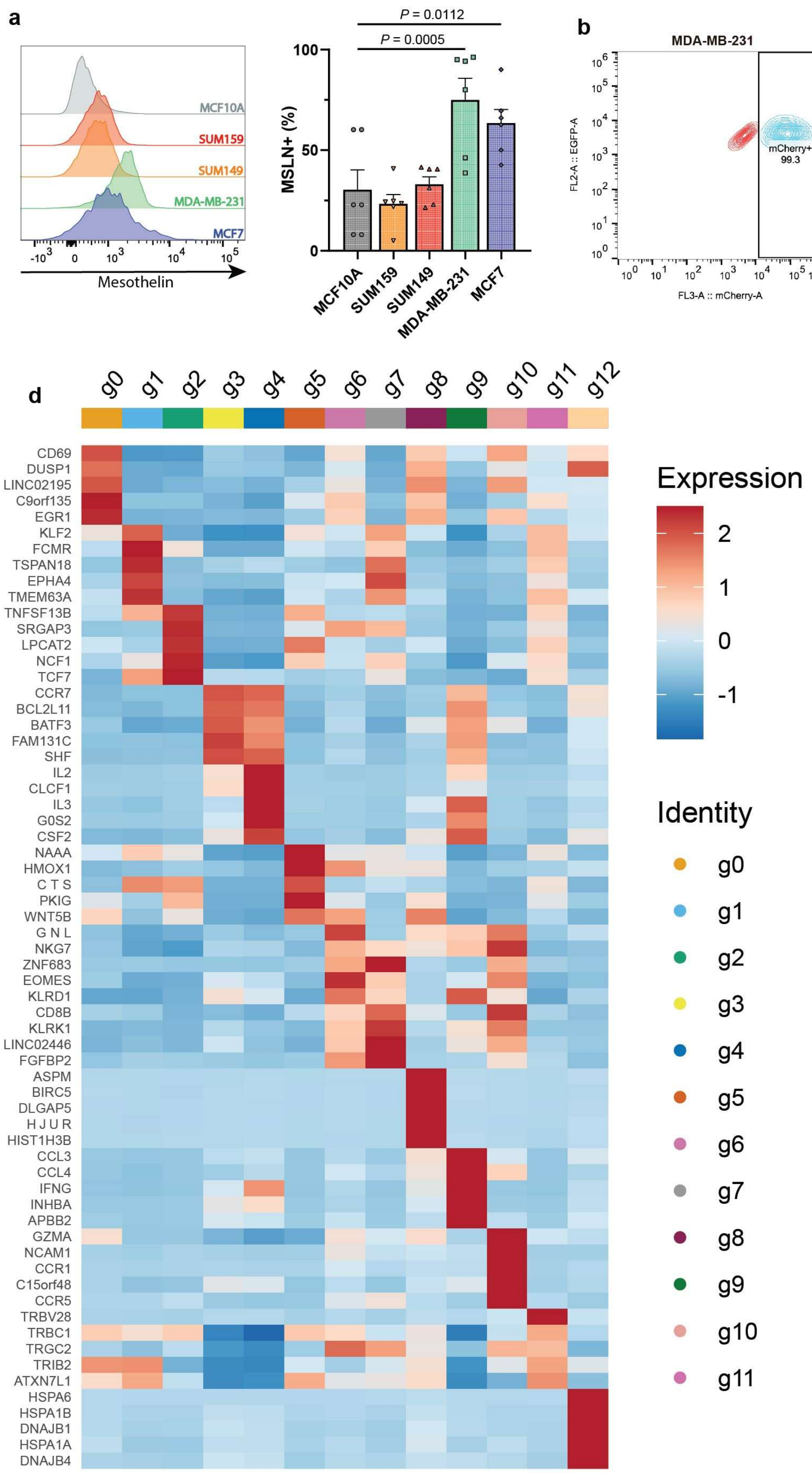
a
MCF10A
SUM159
SUM149
MDA-MB-231
MCF7
Mesothelin
P = 0.0112
P = 0.0005
MSLN+ (%)
100
50
0
MCF10A
SUM159
SUM149
MDA-MB-231
MCF7
b
MDA-MB-231
FL2-A :: EGFP-A
FL3-A :: mCherry-A
mCherry+
99.3
c
MCF-7
FL2-A :: EGFP-A
FL3-A :: mCherry-A
mCherry+
51.7
d
g0
g1
g2
g3
g4
g5
g6
g7
g8
g9
g10
g11
g12
CD69
DUSP1
LINC02195
C9orf135
EGR1
KLF2
FCMR
TSPAN18
EPHA4
TMEM63A
TNFSF13B
SRGAP3
LPCAT2
NCF1
TCF7
CCR7
BCL2L11
BATF3
FAM131C
SHF
IL2
CLCF1
IL3
G0S2
CSF2
NAAA
HMOX1
C T S
PKIG
WNT5B
G N L
NKG7
ZNF683
EOMES
KLRD1
CD8B
KLRK1
LINC02446
FGFBP2
ASPM
BIRC5
DLGAP5
H J U R
HIST1H3B
CCL3
CCL4
IFNG
INHBA
APBB2
GZMA
NCAM1
CCR1
C15orf48
CCR5
TRBV28
TRBC1
TRGC2
TRIB2
ATXN7L1
HSPA6
HSPA1B
DNAJB1
HSPA1A
DNAJB4
Expression
2
1
0
-1
Identity
g0
g1
g2
g3
g4
g5
g6
g7
g8
g9
g10
g11

**Supplementary Figure 4. Validation of BC cell lines and CAR T.** (a) HR+ and TNBC cell lines were assessed for mesothelin expression and compared against MCF10A (an immortalized breast epithelial cell line). N = 2 independent experiments; n = 3 technical replicates. (b) Expression of mCherry in luciferase-mCherry-transduced MCF-7 and (c) MDA-MB-231 cell lines. (c) Expression of the top5 marker genes per UMAP cluster. Columns represent individual clusters and rows represent genes. Color intensity indicates relative expression level (row-scaled z-score).

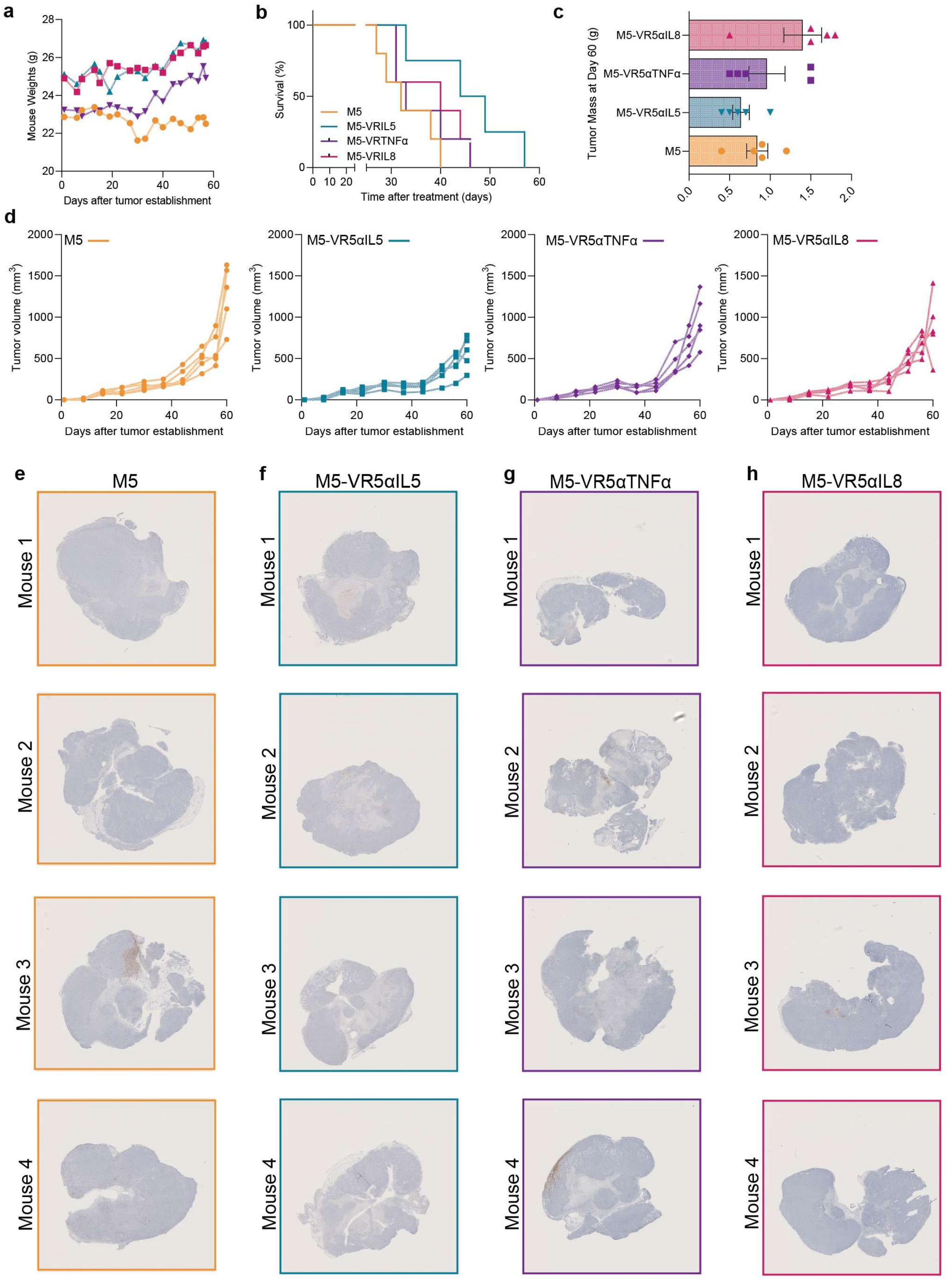
a
Mouse Weights (g)
Days after tumor establishment
b
Survival (%)
M5
M5-VRIL5
M5-VRTNFα
M5-VRIL8
Time after treatment (days)
c
Tumor Mass at Day 60 (g)
M5-VR5αIL8
M5-VR5αTNFα
M5-VR5αIL5
M5
d
M5
M5-VR5αIL5
M5-VR5αTNFα
M5-VR5αIL8
Tumor volume (mm3)
Days after tumor establishment
e
M5
f
M5-VR5αIL5
g
M5-VR5αTNFα
h
M5-VR5αIL8
Mouse 1
Mouse 2
Mouse 3
Mouse 4

**Supplementary Figure 5. VR-CAR-T cells reduce tumor burden.** (a) Mouse weights of each treatment groups monitored over the course of the study. (b) Kaplan-Meier survival curves showing percent survival over time after treatment. (c) Tumor mass at end the experiment (day 60). Statistical significance was assessed by one-way ANOVA with Tukey's multiple comparisons test (*P < 0.05, **P < 0.01, ***P < 0.001, ****P < 0.0001). (d) individual tumor growth curves for each treatment group following tumor establishment. (a-d) n = 5 miceper group. (e-h) Human CD45 IHC staining of tumor sections to identify CAR-T infiltraiton. n = 4 mice per group.

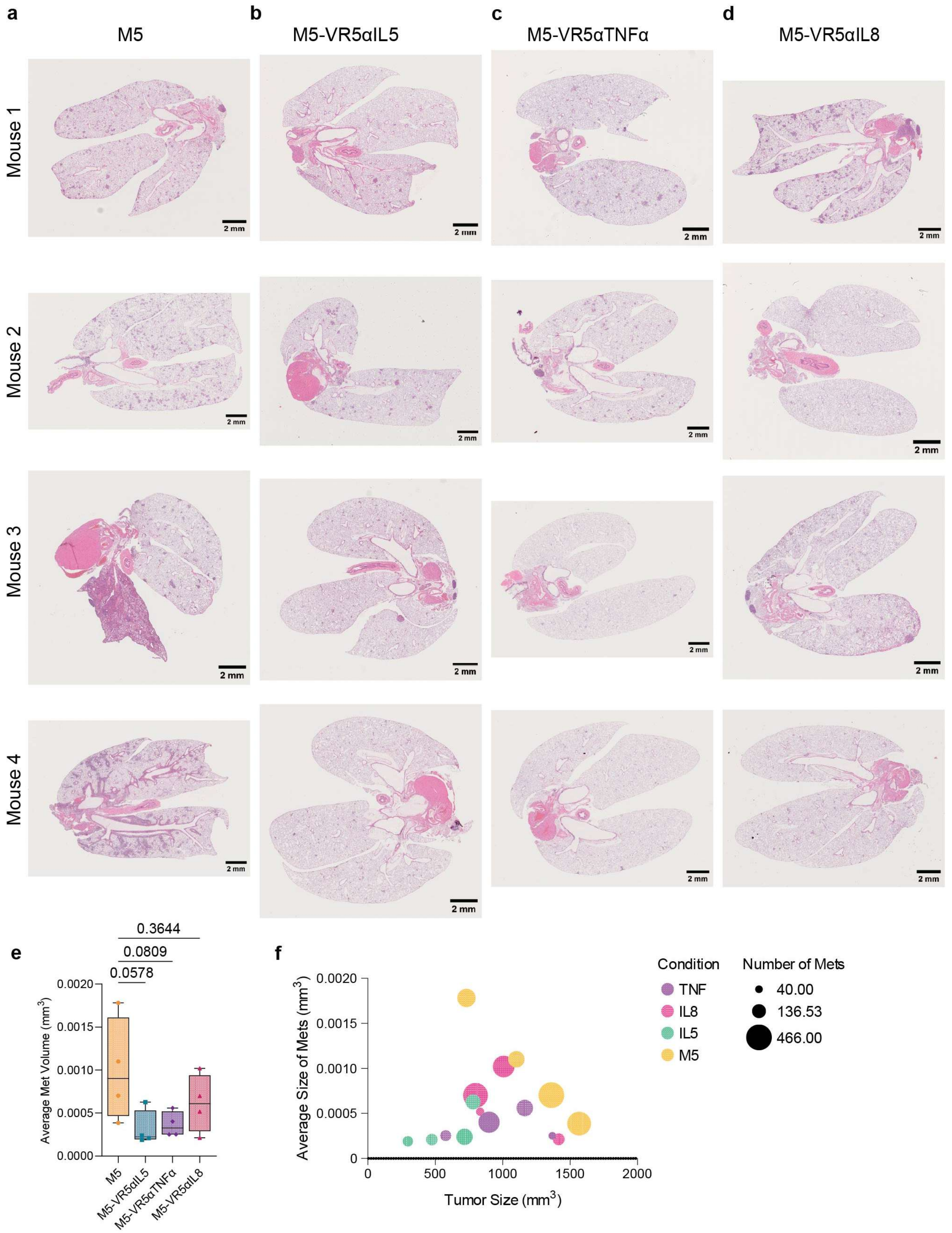
a
b
c
d
M5
M5-VR5αIL5
M5-VR5αTNFα
M5-VR5αIL8
Mouse 1
Mouse 2
Mouse 3
Mouse 4
2 mm
e
0.3644
0.0809
0.0578
Average Met Volume (mm3)
0.0020
0.0015
0.0010
0.0005
0.0000
M5
M5-VR5αIL5
M5-VR5αTNFα
M5-VR5αIL8
f
Average Size of Mets (mm3)
Tumor Size (mm3)
0
500
1000
1500
2000
Condition
TNF
IL8
IL5
M5
Number of Mets
40.00
136.53
466.00

**Supplementary Figure 6. VR-CAR-T cells inhibit lung metastasis.** (a-d) Lung H&E images showing reduced metastatic burden in VR-CAR-T cell-treated mice compared to the M5 control. Scale bar = 2mm. (e) Quantification of the average metastatic volume. Data plotted as boxplots, n = 4 mice per group. One-way ANOVA with Tukey's multiple comparison test was used for statistical significance. (*$P<0.05$, **$P<0.01$, ***$P<0.001$, $P<0.0001$). (f) Bubble plot correlates average metastatic lesion size and number of metastatic lesions with tumor size for each condition.

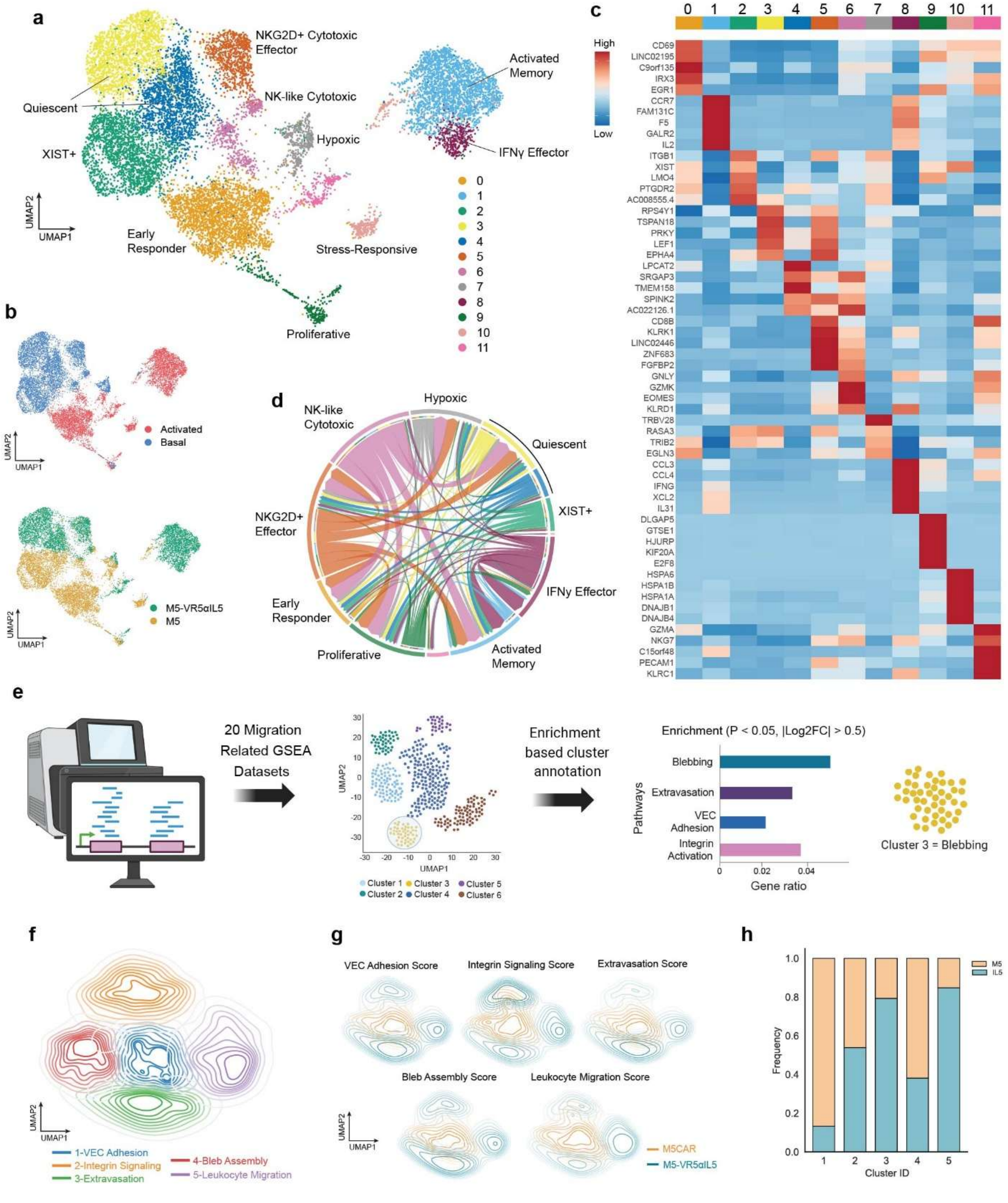


**Supplementary Figure 7.** (a) Unsupervised UMAP embedding of mesothelin-stimulated CAR T cells, colored by transcriptionally defined clusters. (b) Same embedding stratified by activation state and experimental condition. (c) Heatmap of the top-ranked differentially expressed genes

(DEGs) defining each cluster in (a). (d) Cell–cell communication network inferred by CellChat analysis in VR5αIL5-M5 CAR T cells, revealing extensive signaling interactions among activated and cytotoxic subsets. (e) UMAP embedding constructed using genes from 20 curated migration-related gene sets (GSEA; Supplementary Table X), with clusters annotated by highest gene set enrichment. (f) Density contour projection of cells across the migration-focused UMAP space. (g) Condition-resolved contour overlays highlighting enrichment of VR5αIL5 CAR T cells within extravasation- and leukocyte migration-associated regions. (h) Quantification of VR5αIL5 CAR T cell frequency across clusters defined in (e).

**Supplementary Table 1.** GSEA datasets used in migration pathway analysis

| Dataset | # of Genes | URL |
|---|---|---|
| GOBP_CELL_MIGRATION | 1557 | https://www.gsea-msigdb.org/gsea/msigdb/human/download_geneset.jsp?geneSetName=GOBP_CELL_MIGRATION&fileType=grp |
| GOBP_CELL_MOTILITY | 1863 | https://www.gsea-msigdb.org/gsea/msigdb/mouse/download_geneset.jsp?geneSetName=GOBP_CELL_MOTILITY&fileType=grp |
| GOBP_ACTIN_FILAMENT_BASED_PROCESS | 834 | https://www.gsea-msigdb.org/gsea/msigdb/human/download_geneset.jsp?geneSetName=GOBP_ACTIN_FILAMENT_BASED_PROCESS&fileType=grp |
| GOBP_BLEB_ASSEMBLY | 11 | https://www.gsea-msigdb.org/gsea/msigdb/human/download_geneset.jsp?geneSetName=GOBP_BLEB_ASSEMBLY&fileType=grp |
| GOBP_FOCAL_ADHESION_ASSEMBLY | 85 | https://www.gsea-msigdb.org/gsea/msigdb/human/download_geneset.jsp?geneSetName=GOBP_FOCAL_ADHESION_ASSEMBLY&fileType=grp |
| GOBP_INTERMEDIATE_FILAMENT_BASED_PROCESS | 95 | https://www.gsea-msigdb.org/gsea/msigdb/human/download_geneset.jsp?geneSetName=GOBP_INTERMEDIATE_FILAMENT_BASED_PROCESS&fileType=grp |
| GOBP_INTEGRIN_ACTIVATION | 26 | https://www.gsea-msigdb.org/gsea/msigdb/human/download_geneset.jsp?geneSetName=GOBP_INTEGRIN_ACTIVATION&fileType=grp |
| GOBP_INTEGRIN_MEDIATED_SIGNALING_PATHWAY | 113 | https://www.gsea-msigdb.org/gsea/msigdb/human/download_geneset.jsp?geneSetName=GOBP_INTEGRIN_MEDIATED_SIGNALING_PATHWAY&fileType=grp |
| GOBP_POSITIVE_REGULATION_OF_INTEGRIN_ACTIVATION | 11 | https://www.gsea-msigdb.org/gsea/msigdb/human/download_geneset.jsp?geneSetName=GOBP_POSITIVE_REGULATION_OF_INTEGRIN_ACTIVATION&fileType=grp |

| | | |
|---|---|---|
| GOBP_REGULATION_OF_INTEGRIN_ACTIVATION | 17 | https://www.gsea-msigdb.org/gsea/msigdb/human/download_geneset.jsp?geneSetName=GOBP_REGULATION_OF_INTEGRIN_ACTIVATION&fileType=grp |
| GOBP_CELLULAR_EXTRAVASATION | 75 | https://www.gsea-msigdb.org/gsea/msigdb/human/download_geneset.jsp?geneSetName=GOBP_CELLULAR_EXTRAVASATION&fileType=grp |
| GOBP_LEUKOCYTE_ADHESION_TO_ARTERIAL_ENDOTHELIAL_CELL | 6 | https://www.gsea-msigdb.org/gsea/msigdb/human/download_geneset.jsp?geneSetName=GOBP_LEUKOCYTE_ADHESION_TO_ARTERIAL_ENDOTHELIAL_CELL&fileType=grp |
| GOBP_LEUKOCYTE_ADHESION_TO_VASCULAR_ENDOTHELIAL_CELL | 58 | https://www.gsea-msigdb.org/gsea/msigdb/human/download_geneset.jsp?geneSetName=GOBP_LEUKOCYTE_ADHESION_TO_VASCULAR_ENDOTHELIAL_CELL&fileType=grp |
| GOBP_LEUKOCYTE_CHEMOTAXIS | 237 | https://www.gsea-msigdb.org/gsea/msigdb/human/download_geneset.jsp?geneSetName=GOBP_LEUKOCYTE_CHEMOTAXIS&fileType=grp |
| GOBP_LEUKOCYTE_MIGRATION | 399 | https://www.gsea-msigdb.org/gsea/msigdb/human/download_geneset.jsp?geneSetName=GOBP_LEUKOCYTE_MIGRATION&fileType=grp |
| GOBP_REGULATION_OF_LEUKOCYTE_MIGRATION | 232 | https://www.gsea-msigdb.org/gsea/msigdb/human/download_geneset.jsp?geneSetName=GOBP_REGULATION_OF_LEUKOCYTE_MIGRATION&fileType=grp |
| GOBP_REGULATION_OF_LEUKOCYTE_TETHERING_OR_ROLLING | 18 | https://www.gsea-msigdb.org/gsea/msigdb/human/download_geneset.jsp?geneSetName=GOBP_REGULATION_OF_LEUKOCYTE_TETHERING_OR_ROLLING&fileType=grp |
| GOBP_NUCLEUS_ORGANIZATION | 150 | https://www.gsea-msigdb.org/gsea/msigdb/human/download_geneset.jsp?geneSetName=GOBP_NUCLEUS_ORGANIZATION&fileType=grp |

| | | |
|---|---|---|
| GOBP_NUCLEUS_LOCALIZATION | 32 | https://www.gsea-msigdb.org/gsea/msigdb/human/download_geneset.jsp?geneSetName=GOBP_NUCLEUS_LOCALIZATION&fileType=grp |
| GOBP_ARP2_3_COMPLEX_MEDIATED_ACTIN_NUCLEATION | 46 | https://www.gsea-msigdb.org/gsea/msigdb/human/download_geneset.jsp?geneSetName=GOBP_ARP2_3_COMPLEX_MEDIATED_ACTIN_NUCLEATION&fileType=grp |

**Supplementary Table 2.** Definition of features used in multidimensional motility analysis

| *Motility feature* | *Feature type* | *Equation* |
|---|---|---|
| Total distance | Displacement | Sum of instantaneous Euclidean distances in a trajectory |
| Average speed | Displacement | Mean instantaneous Euclidean distance covered in a trajectory |
| Max speed | Displacement | Maximum speed achieved within a trajectory |
| Min speed | Displacement | Minimum speed achieved within a trajectory |
| Net distance | Displacement | Euclidean distance between a cell's position at t=0 and t=59 mins |
| MSD1 | Displacement | Sum of squared distances with time lag=1 |
| MSD2 | Displacement | Sum of squared distances with time lag=2 |
| MSD3 | Displacement | Sum of squared distances with time lag=3 |
| Alpha | Diffusivity | Slope of the log(time lag) vs log(MSD) |
| Progressivity | Persistence | Ratio of the net distance traveled to the total distance traveled |
| Average angle | Persistence | Average turning angle of a cell within a trajectory |
| Min angle | Persistence | Minimum turning angle of a cell within a trajectory |
| Persistence (for microchannel confinement) | Persistence | Ratio of cell velocity to cell speed |
| Displ. variance | Distribution | Variance of displacement distribution within a trajectory |
| Displ. covariance | Distribution | Ratio of displacement standard deviation over mean displacement |
| Displ. skewness | Distribution | Skewness of displacement distribution within a trajectory |
| Displ. kurtosis | Distribution | Kurtosis of displacement distribution within a trajectory |
| Displ. n-gaussian-alpha | Distribution | Mean of $(\text{displacement})^4 / (3 \times (\text{displacement})^2)^2 - 1$ |
| Displ. gini | Distribution | Gini coefficient of displacement |
| Angle variance | Distribution | Variance of turning angle distribution within a trajectory |
| Angle covariance | Distribution | Ratio of turning angle standard deviation over mean turning angle |
| Angle skewness | Distribution | Skewness of turning angle distribution within a trajectory |
| Angle kurtosis | Distribution | Kurtosis of turning angle distribution within a trajectory |
| Angle n-gaussian-alpha | Distribution | Mean of $(\text{angle})^4 / (3 \times (\text{angle})^2)^2 - 1$ |
| Angle gini | Distribution | Gini coefficient of turning angle |
| Displ. autocorr. 1 | Temporal correlation | Autocorrelation coefficient for displacement with time lag=1 |
| Displ. autocorr. 2 | Temporal correlation | Autocorrelation coefficient for displacement with time lag=2 |
| Displ. autocorr. 3 | Temporal correlation | Autocorrelation coefficient for displacement with time lag=3 |
| Displ. partial autocorr. 2 | Temporal correlation | Partial autocorrelation coefficient for displacement with time lag=2 |
| Displ. partial autocorr. 3 | Temporal correlation | Partial autocorrelation coefficient for displacement with time lag=3 |
| Angle autocorr. 1 | Temporal correlation | Autocorrelation coefficient for turning angle with time lag=1 |
| Angle autocorr. 2 | Temporal correlation | Autocorrelation coefficient for turning angle with time lag=2 |
| Angle autocorr. 3 | Temporal correlation | Autocorrelation coefficient for turning angle with time lag=3 |
| Angle partial autocorr. 2 | Temporal correlation | Partial autocorrelation coefficient for turning angle with time lag=2 |
